# Therapeutic targeting of oligodendrocytes in an agent-based model of multiple sclerosis

**DOI:** 10.1101/2025.06.26.661705

**Authors:** Georgia R Weatherley, Robyn P Araujo, Samantha J Dando, Adrianne L Jenner

## Abstract

Multiple sclerosis (MS) is a neurodegenerative disease in which misdirected, persistent activity of the immune system degrades the protective myelin sheaths of nerve axons. Historically, treatment of MS has relied on disease-modifying therapies that involve immunosuppression, such as targeting of the blood-brain barrier (BBB) to restrict lymphocyte movement. New therapeutic ideas in the development pipeline are instead designed to promote populations of myelin producing cells, oligodendrocytes, by exploiting their innate resilience to the stressors of MS or restoring their numbers. Given the significant advancements made in immunological disease understanding due to mathematical and computational modelling, we sought to develop a platform to (1) interrogate our understanding of the neuroimmunological mechanisms driving MS development and (2) examine the impact of different therapeutic strategies. To this end we propose a novel, open-source, agent-based model of lesion development in the CNS. Our model includes crucial populations of T cells, perivascular macrophages, and oligodendrocytes. We examine the sensitivity of the model to key parameters related to disease targets and conclude that lesion stabilisation can be achieved when targeting the integrated stress response of oligodendrocytes. Most significantly, complete prevention of lesion formation is observed when a combination of approved BBB-permeability targeting therapies and integrated-stress response targeting therapies is administered, suggesting the potential to strike a balance between a patient’s immune inflammation and their reparative capacity. Given that there are many open questions surrounding the etiology and treatment of MS, we hope that this malleable platform serves as a tool to test and generate further hypotheses regarding this disease.

**Author summary:** Multiple sclerosis is a disease that is not yet fully understood and has no cure. Some patient phenotypes see little benefit from current therapeutic interventions besides symptomatic treatment. Typically, MS studies have focused on the prevention of damage to brain tissue. As such, there are unanswered questions about how to reverse the damage in the brain and spinal cord of MS patients caused by immune cells. In light of this there is an urgent need for mathematical modelling of new therapeutic strategies - some of which remain to be clinically examined - shifting the attention from the targeting of aberrant immune activity to the upregulation of resident, reparative cells called oligodendrocytes. Here, we have developed a mathematical model to probe the potential benefits of oligodendrocyte targeted therapies *in silico*. We focus on T cells as damaging agents and monitor oligodendrocyte function in response to their activity. We show that oligodendrocytes strongly influence the tissue’s ability to stabilise and even recover under persistent, harmful immune activity. In practice, these therapies could hold the potential to unlock neuroprotection by means of enhanced remyelination.

## Introduction

Multiple sclerosis (MS) is a neuroimmune disease with increasing global prevalence and disproportionate incidence in women [1]. It is pathologically hallmarked by inflammation and neurodegeneration, whereby demyelination and axonal loss give rise to lesion formation and impaired central nervous system (CNS) function (see Fig 1) [2, 3]. While the exact disease etiology is unclear, MS is thought to be induced by the interaction of inherent genetic susceptibility with environmental and lifestyle risks that include viral infections, low vitamin D levels and smoking [4–6]. The complexity of these interactions gives rises to highly heterogeneous disease courses with notable variations between treatment responses and clinical outcomes of different patient cohorts, preventing a uniform approach to treatment [7, 8].

**Fig 1.**
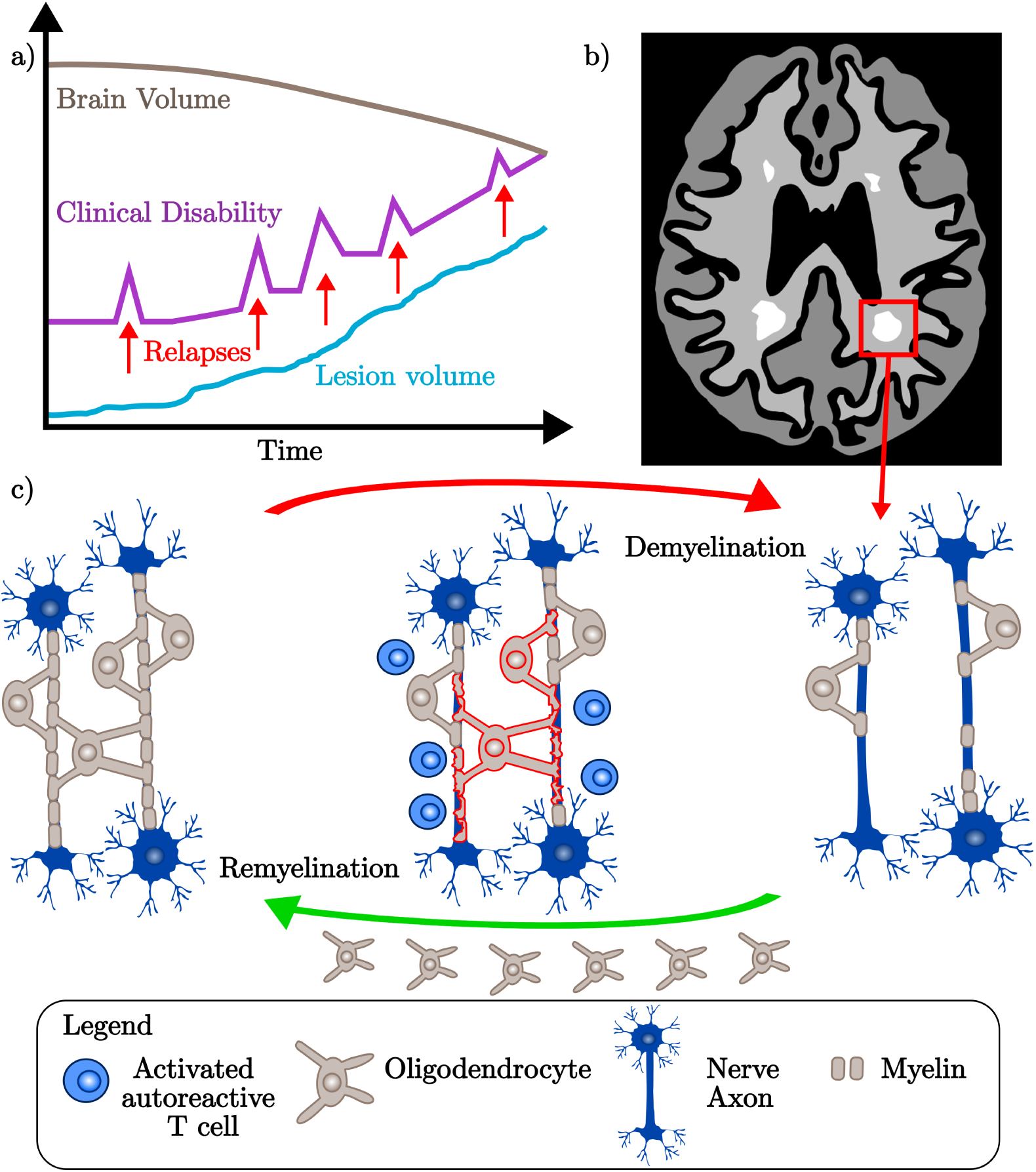
**a** When MS patients experience relapses, there is a spike in their clinical disability. In addition to these inflammatory events, the underlying accumulation of permanent damage results in decreased brain volume as there is a gradual increase of lesion volume (subclinical disease). **b** MRI scans serve as both a diagnostic tool and an ongoing measure of disease progression. Here we show a cartoon map depicting a typical axial slice of a patient’s MRI data. Lesions are shown as white areas of demyelination. **c** In MS, the demyelination of nerve axons is driven by immune cells such as activated T cells. Myelin is a membrane extension of oligodendrocytes, which are cells that undertake remyelination to maintain proper nerve signalling. Myelin loss in MS patients removes the protective insulation of nerve cells, causing oligodendrocyte loss and symptom onset subject to lesion location (recall **b**).

The damage associated with MS arises from the complex interplay of both the innate and adaptive immune responses (Fig 1c) [9]. Activated lymphocytes, microglia and macrophages are the primary drivers of the sustained, misdirected attack on myelin sheaths that surround, insulate, and protect nerve axons [9]. The lymphocytes, such as T cells and B cells, are activated in the periphery and cross the blood-brain barrier (BBB) to infiltrate the CNS [10]. A functional BBB regulates movement into the CNS, limiting the intake of toxins and immune cells. Dysregulation of the BBB in MS patients leads to increased lymphocyte migration, heightening the local myelin loss (demyelination) of nerve axons [11]. This causes areas of scarring called lesions, signifying demyelination has occurred. While we attribute lesion development to lymphocyte activity, incomplete lesion recovery (remyelination) is attributed to oligodendrocyte loss [12]. Oligodendrocytes are myelin synthesising cells of the CNS that sustain myelin rates over the long term for effective neuronal signalling [13]. The inflammation associated with MS is cytotoxic to oligodendrocytes, with their loss furthering disease pathogenesis.

The different phenotypes of MS (Fig 1a) are distinguished by relative differences in disease activity and the extent of accumulated damage. It is estimated that 85% of patients initially follow a relapsing disease course, with the remaining 15% of patients considered primary progressive (PPMS) [14–16]. Relapsing-remitting MS (RRMS) patients experience intermittent clinical relapses that are driven by heightened lesion activity and neural disruption [17]. Relapses are marked by the rise of new symptoms or exacerbations of previous symptoms. In primary progressive (PPMS) and secondary progressive (SPMS) patients, the dominant driver of disability is the accumulation of progressive damage rather than the large, active inflammatory events experienced by RRMS patients [5]. Over the course of the disease, this causes decreases in brain volume. Diagnosis of MS and classification of the patient’s phenotype is conducted using magnetic resonance imaging (MRI), see Fig 1b.

Current medical interventions aim to improve patient wellbeing through the prescription of disease-modifying therapies (DMTs) and symptomatic treatments [18]. Unfortunately, there is no cure for MS and disease phenotype has a significant impact on a patient’s access to disease-altering treatments [19]. While the first DMT for RRMS patients was approved in 1993 (IFNβ-1b), the first DMT for PPMS patients was only approved in 2018 (ocrelizumab) [20, 21]. DMTs are designed to reduce relapse frequency and inflammatory activity [3]. They often rely on blocking lymphocyte infiltration into the CNS, such as natalizumab which disrupts cell adhesion through the integrin α4 [22]. Over the long term, these immunosuppressant therapies have been recorded to increase the patient’s risk of complications arising from viral infections in the brain [23]. Further, DMTs are limited in their ability to reduce a patient’s existing functional deficits. While there are ongoing efforts to identify further therapies that target the T cells that mediate MS [24, 25], there is also recent interest in new therapies promoting remyelination [3]. It is anticipated that these new remyelination therapies would restore myelination within lesions by promoting increased oligodendrocyte resilience or remyelination rates [26].

There are several types of remyelination therapies being proposed. Those relying on increased protection of mature oligodendrocytes aim to enhance their innate integrated stress response (ISR) to suppress apoptosis under the stresses of MS [27]. Cellular treatments instead aim to achieve remyelination by introducing new oligodendrocytes into demyelinated areas using stem cell treatments or oligodendrocyte precursor cells (OPCs) [28, 29]. OPCs are resident to the CNS and can differentiate into mature oligodendrocytes to promote new myelin [30]. PPMS and SPMS patients stand to benefit significantly from these new therapeutic ideas given their large accumulated losses of myelin. There is even suggestion that the clinical outcomes of these progressive patients may be improved by the combined use of immunosuppressive therapies (DMTs) and remyelination therapies [30].

The aspects of MS that make it a challenging disease to predict and treat persist when modelling it mathematically. Nonetheless, both stochastic and deterministic models of MS have been used to advance our understanding of the underlying drivers of the different disease courses, and refine potential therapeutic strategies for their treatment (see review in Weatherley *et al.* [31]). Partial differential equation (PDE) systems have largely focused on understanding patterns of lesion formation [32–34], with the work of Moise and Friedman [35] being the first PDE model to relate lesion volume to the timing and combinations of established and experimental DMTs. While models such as these are effective at capturing myelin loss at the scale of the lesion they are unable to account for the inherent heterogeneity of MS. This motivates interest in stochastic models that can still capture the dissemination of MS in both space and time [36].

Agent-based models (ABMs) have been used to investigate drivers of neurological disease, such as the spatiotemporal distribution of atrophy in Parkinson’s disease [37] and the roles of the olfactory system [38] and microglia [39] in Alzheimer’s disease. However, there are few open-source ABMs that model the key cell-cell interactions of MS [40–44]. Using the Netlogo framework, Pennisi *et al*. [40] simulated a 51 by 51 cell grid of brain and blood regions to explore BBB-specific therapeutic interventions. The model incorporates various immune cell populations and myelin alongside a dynamic BBB designed to mimic the BBB behaviour of MS patients, ultimately suggesting that there is greater treatment effectiveness in preventing BBB leakage rather than attempting recovery of BBB functionality. The model built upon previous ABMs that examine the cross-regulatory behaviours of regulatory T cells and effector T cells [41] and examine the potential of Vitamin D in MS treatment given its immunomodulatory ability [42]. This body of work has contributed to the development of the Universal Immune System Simulator (UISS). In addition to simulating tuberculosis [45], carcinoma [46], and COVID-19 [47], this software has recently been used to predict the outcome of simulated RRMS patients treated with DMTs [43, 44]. A lack of open-source access to these existing models hinders our ability to build upon this work.

Here, we develop an ABM to investigate the potential of oligodendrocytes as novel therapeutic targets in the treatment of MS. It has been suggested that some MS patients may see better outcomes from remyelination treatments concerned with oligodendrocyte protection and the promotion of remyelination, rather than purely with the suppression of harmful immune activity (DMTs) [30]. Further, Broome and Coleman (2011) [48] have modelled cellular pathways of oligodendrocyte apoptosis using biochemical systems theory to highlight their significance as therapeutic targets. Developed in Matlab and with open-source code provided, we believe our on-lattice ABM to be the first in the MS context to incorporate the activity of oligodendrocytes. By understanding the effects of oligodendrocyte loss and oligodendrocyte preservation on myelination rates, we hope to elucidate the significance of these key cells that underpin recently proposed remyelination therapies.

We first outline the methods, with a particular focus on how we capture the underlying intracellular dynamics that determine the outcomes of individual oligodendrocytes. In the Results, we demonstrate the capabilities of the model, and how its design captures both cell-based and lesion-level outcomes. By undertaking a sensitivity analysis to understand the significance of key parameters, we use this insight to highlight how perturbations in oligodendrocyte behaviours mimicking treatment effects could alter the expected disease course. In Table 1 we present the key terminologies and abbreviations used throughout this paper.

**Table 1.**
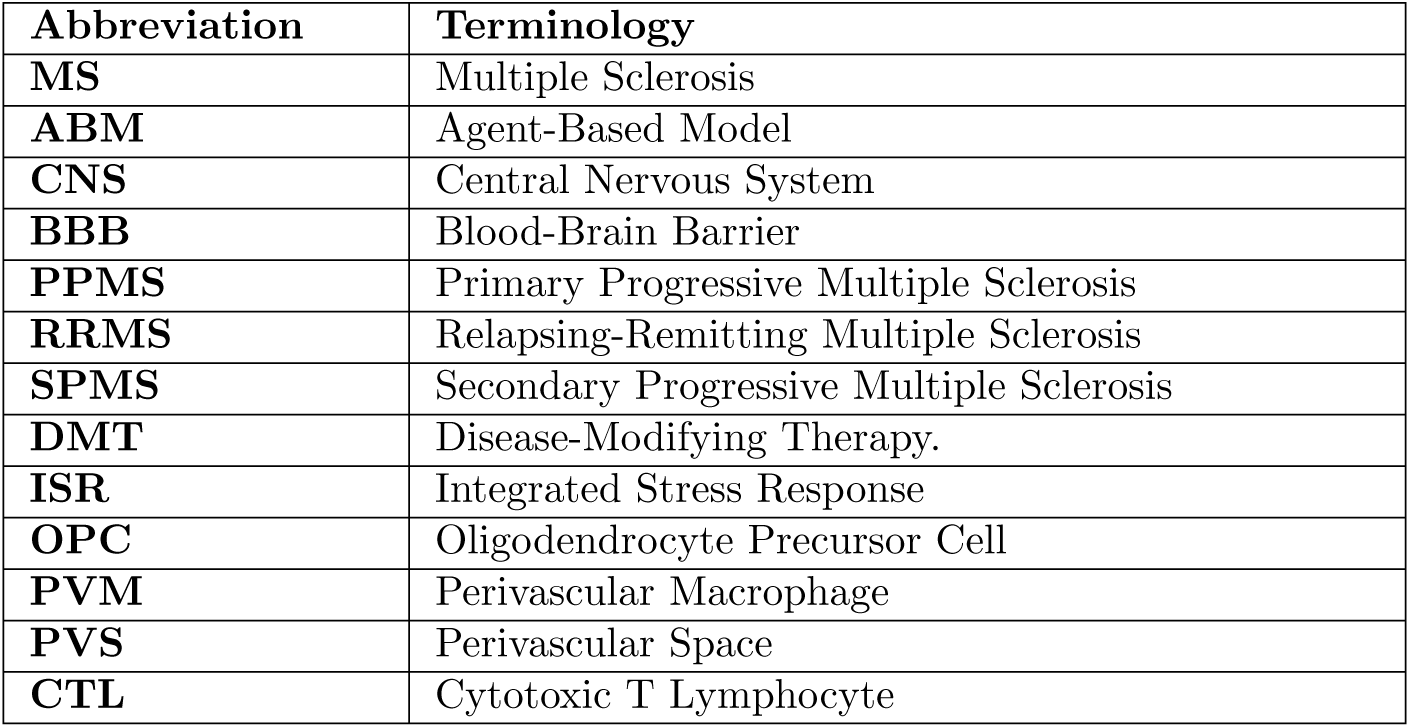
Abbreviations of key terminology.

## Methods

In this model, our focus is on capturing the disruption to typical oligodendrocyte and myelin homeostasis introduced by the aberrant immune activity of MS. We condense immune cell activity to consider only primed T cells, perivascular macrophages (PVMs), and reactivated T cells. As such, the model includes only the following agent populations: myelin, oligodendrocytes, primed T cells, PVMs, and reactivated T cells (see Fig 2). These capture all necessary disease elements to simulate long-term demyelination resulting from harmful lymphocyte activity and reparative oligodendrocyte behaviours. These agent populations undergo events such as cell movement, cell renewal, cell death, myelin repair and myelin destruction. We give further description of these events when outlining each agent and its implementation. This is an on-lattice model that is discrete in both space and time.

**Fig 2.**
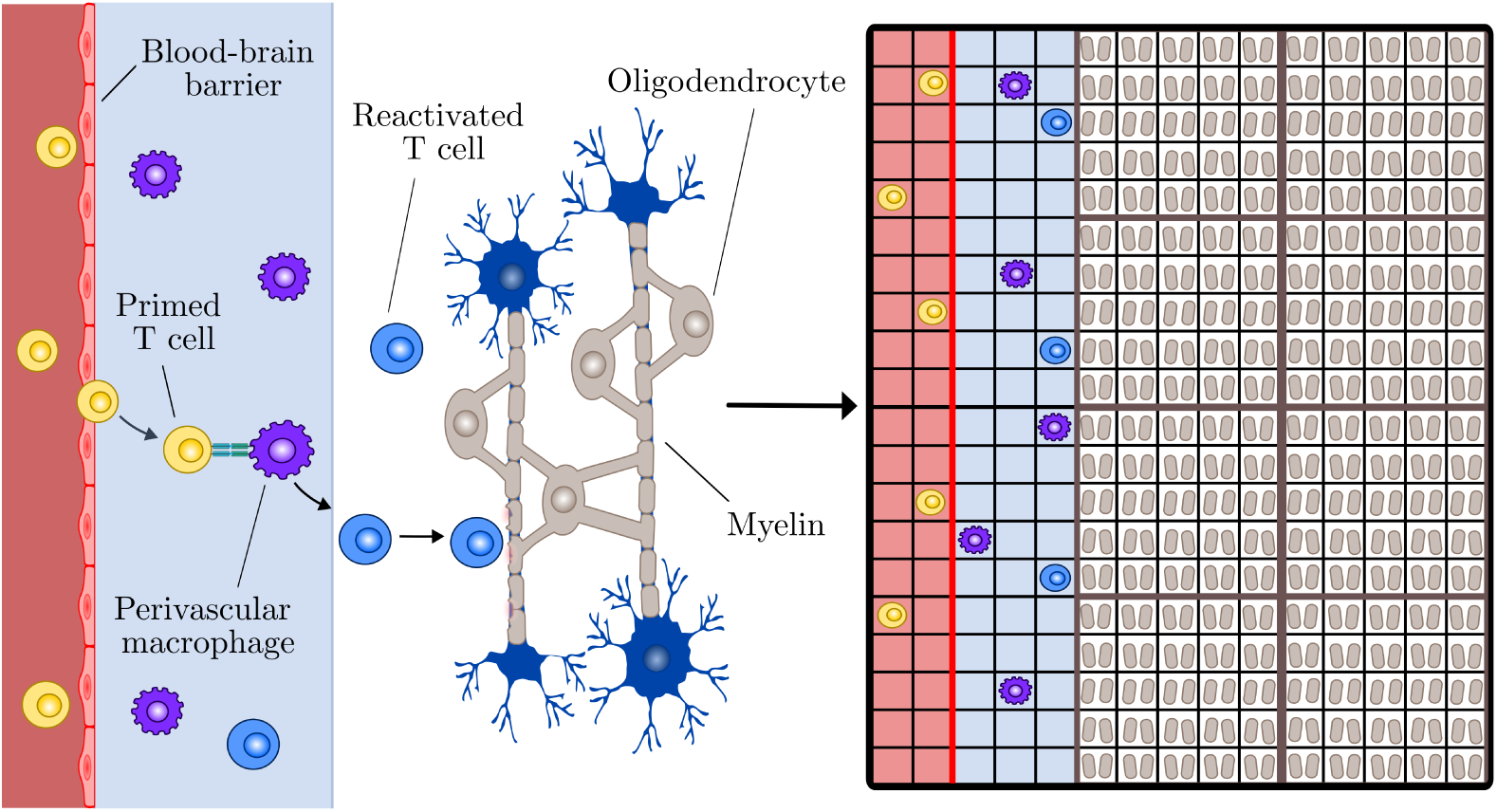
On the left we show a simplification of MS immunology. Following priming in the periphery, autoreactive T cells migrate within blood and then cross the blood-brain barrier to enter the perivascular space. Perivascular macrophages present myelin antigens to T cells resulting in their re-activation and entry into the brain parenchyma. The accumulation of myelin loss and the death of myelin-synthesising cells called oligodendroctyes gives rise to lesion formation. This simplified physiology is translated to a mathematical model, shown on the right (not to scale). A square lattice is divided into regions of peripheral blood (red), perivascular space (blue), and parenchyma (white). Agent populations include primed T cells (yellow), perivascular macrophages (purple), reactivated T cells (blue), myelin (grey) and oligodendrocytes (outlined grey boxes).

### Model domain

We specify a lattice spacing Δ and a time step *τ*. Modelled over a 2-dimensional Cartesian geometry, each immune cell agent is centred on a square lattice site. For instance, a given T cell may occupy position (*x_T_, y_T_*). The immune cell populations are assumed to have uniform diameters of 10*µ*m [49, 50], motivating a uniform lattice spacing of Δ = 10*µ*m. The lattice is 108 units by 300 units and divided into three distinct subdomains: the peripheral blood, the perivascular space (PVS), and parenchyma (see Fig 2). Each region encompasses the full domain height of 300 units, equivalent to 3000*µ*m. We assign a width of 3 units (30*µ*m) to the peripheral blood [51] and a width of 5 units (50*µ*m) to the PVS [35], assuming an enlarged perivascular space under the neuroinflammation of MS [52]. Assigning a width of 100 units (1000*µ*m) to the parenchyma, we model a 3mm^2^ myelinated area along a 3mm major axis. Small MS lesions are typically at least 3mm (3000*µ*m) along their major axis [53]. All parameter choices and their relevant literature sources are summarised in the supplementary material (S1 Full parameter table).

### Relapses

The duration of one time step is *τ* = 20 minutes (see S1 Full parameter table). We advance each simulation over 21,600 time steps to model a 300 day period. While relapse frequency is subject to environmental factors, disease duration, and a patient’s age, sex, and ethnicity [54], diagnostic criteria suggests that the onset of two distinct relapses are separated by at least 30 days [55]. While relapsing episodes are days to weeks in duration [56], we assume a uniform relapse duration, intensity, and frequency. Across a 300 day window we simulate the onset of a new relapse every 100 days, lasting four weeks each, to give three relapse events (see Fig 3d). These relapse events mimic the periodic exacerbations of inflammation typical of relapsing–remitting disease activity [57] (Fig 1a). Taking a relapse to be driven by increased primed T cell infiltration into the CNS, we simulate the event by increasing the number of primed T cells being initialised in the peripheral blood. During a relapse, each lattice site has a probability of *ρ_R_* of generating a time cell in the time step Delta. In a non-relapse period, this probability decreases to *ρ_NR_*, where *ρ_NR_ << ρ_R_*. In the peripheral blood where primed T cells are initialised, this results in an average introduction of 2.25 new primed T cells per time step during relapses, and 0.18 new primed T cells per time step otherwise.

**Fig 3.**
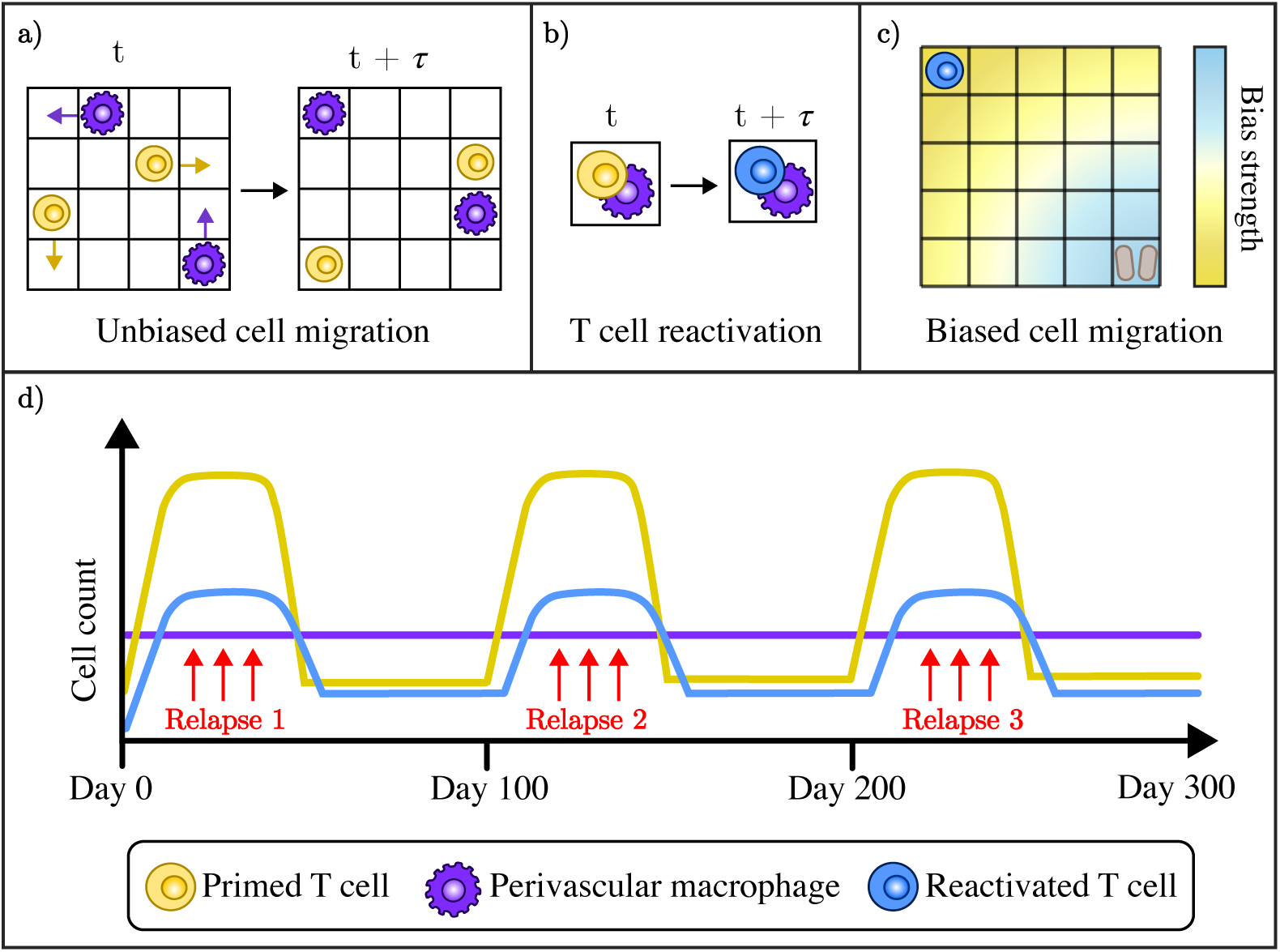
**a** Cells are centred on square lattice sites. Primed T cells (yellow) and PVMs (purple) undertake unbiased migration, with an equal probability of migrating north, south, east or west from their current position to a neighbouring lattice site. **b** The occupation of a lattice site is not mutually exclusive. Same-site occupation of a lattice site by a T cell and a PVM is recorded as activation of that primed T cell, so that it becomes a reactivated T cell (blue), and the PVM remains. **c** The reactivated T cells (blue) are incentivised to undertaken biased random movement towards the nearest myelinated lattice site. The strength of the bias in a reactivated T cell’s movement increases as the cell becomes closer to the myelin (brown). **d** The model is simulated over three hundred days, with three relapse events that are one hundred days apart and four weeks in duration. During each relapse, the inflow of primed T cells is increased, and when these cells enter the PVS there is a delayed increase in the reactivated T cell population.

### Immune cells

There are numerous cell types implicated in the pathogenesis of MS (see [5, 17] for in-depth descriptions of MS immunology). While there is increasing weight attributed to the role of cells such as B cells in the pathogenesis of MS [58], T cells have long been considered to play a central role [59]. Here, we focus on the cellular interactions of T cells and PVMs. Assumed to be circulating in the blood, primed T cell agents migrate across the BBB from the blood region to the PVS. PVMs present myelin antigens to the primed T cell agents, resulting in their reactivation. Reactivated T cell agents have a strictly inflammatory role, programmed to seek and degrade myelin across the parenchyma. The PVM agents solely function as antigen-presenting cells to establish the reactivated T cell presence in the PVS and parenchyma, and we do not account for their ability to undertake phagocytic activity or support the integrity of the BBB [50].

#### Immune agent initialisation

The primed T cell and PVM agents are initialised by randomly generating lattice site coordinates from each of their respective resident regions. For primed T cells this is the peripheral blood, and for PVMs this is the PVS. As in the MS model of Vèlez de Mendizábal [60], we introduce new primed T cells independent to the current primed T cell population density. T cell data has shown their density to vary across lesion location and disease phenotype. We aim to reproduce an average reactivated T cell density of 30-40 cells/mm^2^ in the lesions of RRMS patients [61]. In our model this translates to an average of 110 reactivated T cell agents occupying the 3mm^2^ region of parenchyma during relapse events, and a small, non-zero population in non-relapsing events. We detail our chosen parameters in the supplementary material (S1 Full parameter table).

In a study of PVM distribution subject to brain region, 1-4% of the blood vessel surface was shown to be covered by PVMs [62]. Given limited data availability for PVMs, we estimate a PVM density of 300 cells/mm^2^ in the PVS by assuming 3% occupation of the 0.15mm^2^ area of PVS. Reactivated T cells are only introduced via primed T cell and PVM interaction representative of antigen presentation. This occurs during the simultaneous occupation of a lattice site by a primed T cell agent and a PVM agent (see Fig 3). As implied, cells do not compete for space given lattice site occupation is not mutually exclusive. Following antigen presentation, there is instantaneous removal of the primed T cell agent and introduction of a reactivated T cell agent. The new reactivated T cell agent is initialised at the site of activation, which is still occupied by the PVM. Since PVMs are restricted to their resident domain of the PVS, all events of antigen presentation occur in this region (see Fig 2).

#### T cell death

Further reductions in the primed T cell population arise from death events. The estimated half-life of cytotoxic T lymphocytes (CTLs) is 48 hours, to give a decay rate of 0.35/day [63, 64]. As a result, we define a death probability *ρ_d_* = 0.0049 step for primed T cells and reactivated T cells. Following the successful death event of a primed T cell or reactivated T cell agent where *ρ_d_ > r* ∼ *U* (0, 1), it is immediately removed from the simulation. Given these primed T cell death events, restoring the rate of primed T cell introduction to *ρ_NR_* at the end of each relapse reduces the cell population back to its baseline density.

Expectations of the longevity of PVMs are varied. Some macrophage populations, such as those in the lungs, have been shown to have a 13% turnover after 3.5 years [65]. Further, early studies of rat brain perivascular cells have shown a cell longevity exceeding two years [66]. Simulating across 300 days, we assume a stable cell population of PVMs restricted to the PVS that is not subject to renewal, death or boundary loss.

#### Immune cell movement

All three of the immune cell populations have the capacity to undertake unbiased random walks within a Von Neumann neighbourhood of four neighbouring sites (north, south, east or west on the Cartesian grid). Primed T cells and PVMs are restricted to Brownian motion, as observed *in vitro* for T cells [67]. We impose a successful movement event for each cell at every time step, so that the probability of migrating to a given neighbouring site is 0.25 (see Fig 3).

To recognise the role of chemoattractants such as chemokines, we implement biased movement of reactivated T cells towards myelin following the first instance of damage recorded in the myelin population. While chemokines are not modelled explicitly, our introduction of bias recognises the upregulation of chemokine receptors in T cells following their activation, allowing them to undertake environmentally guided migration [67, 68]. For the reactivated T cells influenced by chemotaxis, we introduce a concentration gradient where the bias, *s*_bias_, in their movement probabilities strengthens as they approach the nearest myelinated site:

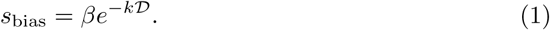

Given *k* is a small, positive constant and D is the Euclidean distance from the reactivated T cell to the nearest myelin agent, the maximum set bias *β* is achieved as the reactivated T cell nears the myelin. *β* is set so that the cell’s movement is not perfectly biased, incorporating the ‘trial and error’ process observed in bacteria and other cells that rely on local relative differences in chemoattractant concentration when determining the biochemical gradient to follow [69].

To consider the directional incorporation of bias, consider a reactivated T cell located at lattice site (*x_T_, y_T_*), where the nearest myelin agent is located south-east at lattice site (*x_j_, y_j_*). For reference, see the described scenario in Fig 3**c**. As *x_T_* − *x_j_ <* 0, and *y_T_* − *y_j_ >* 0, the reactivated T cell is inclined to bias right and down toward the myelin agent. In addition to the standard movement probability of 0.25, the component-wise distribution of the bias *s*_bias_ is determined by the absolute relative magnitude of the *x* and *y* components of D, given by |*x_T_* − *x_j_*| and |*y_T_* − *y_j_*|. For the described problem, the reactivated T cell’s movement probabilities are as follows, where (*x_T_, y_T_*) → (*x_T_* + 1*, y_T_*) represents rightward movement and (*x_T_, y_T_*) → (*x_T_, y_T_* − 1) represents downward movement:

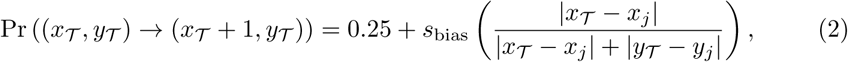

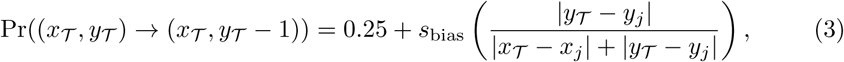

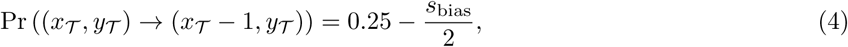

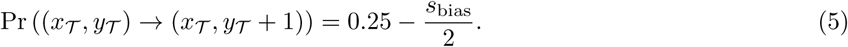

The sum of all movement probabilities is one. See S1 Full parameter table for details on the parameter choices.

### Boundary conditions

As we orientate the peripheral blood vertically to model along the source of incoming lymphocytes (see Fig 2), we enforce periodic (wrap-around) boundary conditions at the top and bottom of the domain. The right boundary has a Dirichlet condition, resulting in the removal of cells if they cross this boundary to travel further into the parenchyma. The left boundary has a Neumann condition, so that the T cells in the peripheral blood can only exit along the blood-brain barrier (BBB) on the right to enter the perivascular space. The role of PVMs as non-parenchymal border-associated macrophages [70] calls for the imposition of an interior boundary between the PVS and parenchyma. When PVMs attempt to enter the parenchyma, we impose a Neumann condition to ensure that they remain in the PVS. Given primed T cells diffuse into the parenchyma from the PVS [61], we allow their unrestricted infiltration across the boundary under a Dirichlet condition.

#### Blood-brain barrier

We incorporate the BBB as an internal boundary that regulates movement between the peripheral blood and perivascular space [71]. The degree of regulation is determined by the BBB permeability level, *b_p_* ∈ [0, 1]. We consider the direction in which immune cells attempt to cross the BBB to define different permeabilities for leftward movement, *b_L_* (*p* = *L*), and rightward movement, *b_R_* (*p* = *R*). Assuming immune cells to be motivated to remain in the CNS, we allow no leftward movement from the CNS to the peripheral blood, achieving a Neumann condition by setting *b_L_* = 0. As BBB breakdown is a hallmark of MS [59, 72], we assume active disruption of BBB functionality from the simulation onset by specifying a non-zero rightward permeability such that *b_R_ >* 0. This allows lymphocyte trafficking into the CNS, whereby an immune agent undergoes a successful movement event across the BBB with probability:

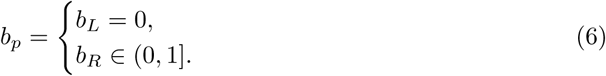

We assume this probability *b_p_* to be a time-invariant, normalised measure of the level of BBB permeability, despite T cell migration compounding BBB disruption in this disease [73].

### Myelin and oligodendrocytes

Unlike the three immune cell populations, the oligodendrocytes and myelin of our model are spatially fixed. We instead track changes in their states and behaviours. Given that myelin is formed as the membrane extension of the oligodendrocyte [74], we closely relate the two populations so that myelin repair is dependent on whether it is associated with a functional oligodendrocyte, and the functionality of each oligodendrocyte is dependent on the extent of damage to its local myelin.

#### Oligodendrocytes

Oligodendrocytes are agents that undertake the repair of myelin following injury [75]. To reconcile the spatial discrepancy between the size of an oligodendrocyte agent and the size of a myelin agent, we assign one myelin agent to each lattice site, with surface coverage of Δ^2^ = 10*µ*m^2^. Each oligodendrocyte agent then uniquely encompasses myelin agents across multiple lattice sites in an approach akin to Cellular Potts. While the surface area of an individual oligodendrocyte’s myelin membrane depends on its type and can reach 50,000 *µ*m^2^ [13], we follow the models of Khonsari and Calvez (2007) [32] and Lombardo *et al*. (2017) [34] to assume an oligodendrocyte density of 400 cells per mm^2^ [7], equivalent to 2500 *µ*m^2^ of myelin membrane per oligodendrocyte. Recalling the lattice spacing of Δ = 10*µ*m, we assume each oligodendrocyte is defined over a 5 × 5 lattice site region to give a total coverage of 2500 *µ*m^2^ across 25 myelinated sites (see Fig 4). Our initial condition establishes a uniform density of myelinating oligodendrocytes in the myelinated region of parenchyma (Fig 2).

**Fig 4.**
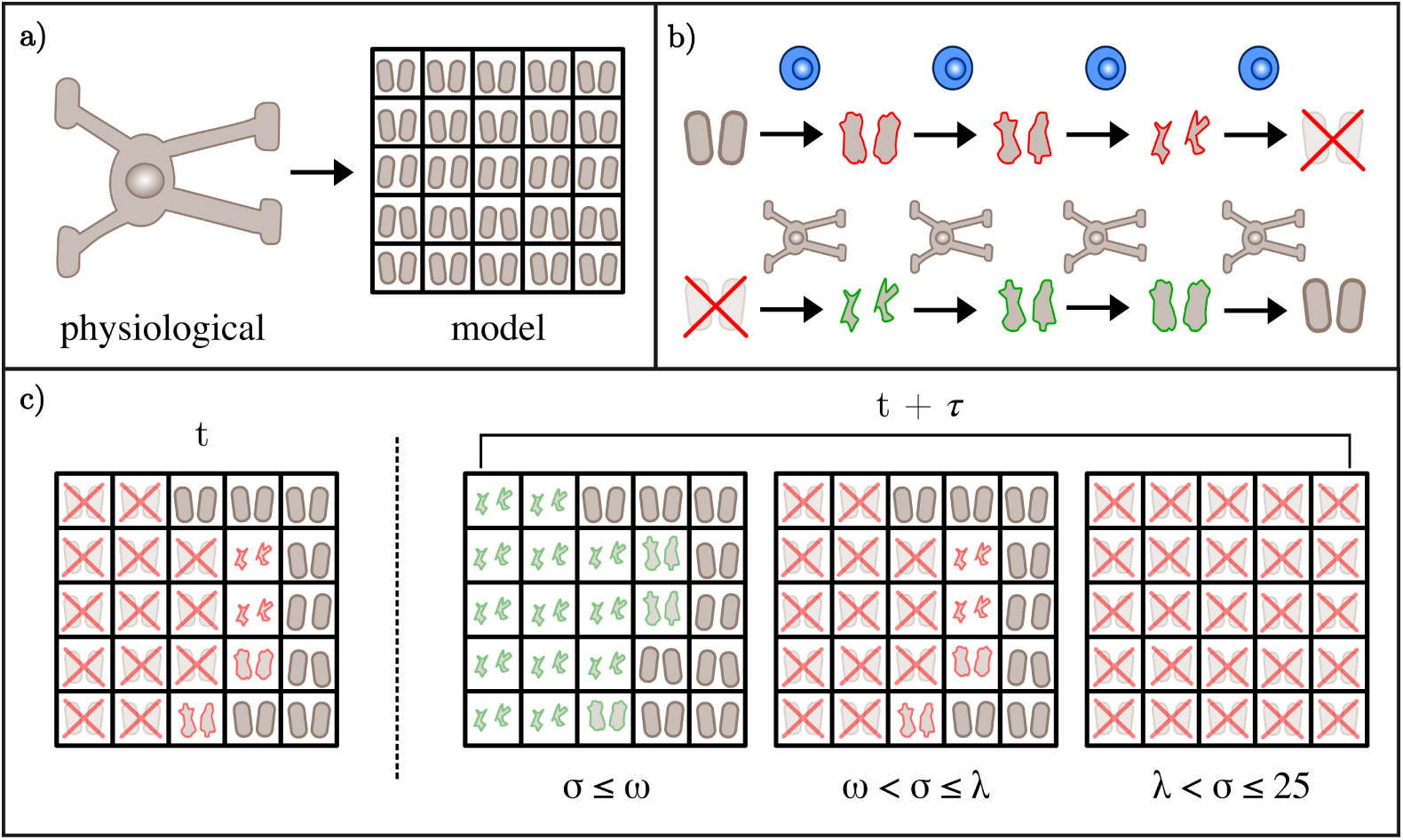
**a** To reconcile the spatial discrepancy between the size of oligodendrocytes and the size of the immune cell populations, we model each oligodendrocyte as a 5 by 5 block of myelinated lattice sites. **b** When a myelinated lattice site is occupied by a reactivated T cell, it is progressively degraded until it is fully demyelinated. If the myelinated lattice site resides in a block associated with an oligodendrocyte that can undertake myelin synthesis, the site is remyelinated over a series of time steps. **c** The extent of demyelination within an oligodendrocyte block determines the extent to which the oligodendrocyte undertakes its typical physiological functions. When the number of completely demyelinated sites *σ* exceeds the first threshold *ω*, the oligodendrocyte loses the ability to remyelinate. Once the total damage *σ* exceed the second threshold *λ*, the oligodendrocyte undergoes apoptosis and all of the associated myelin is removed from its designated myelin sites.

Due to the substantial protein synthesis required for myelin production, oligodendrocytes are sensitive to the disruption of proteostasis [76]. Disruptors implicated in the pathogenesis of MS, such as cytokines, oxidative stress, and nitric oxide, induce cytoprotective mechanisms such as the integrated stress responses (ISR) within oligodendrocytes to promote cell survival [77]. There is strong evidence showing that the ISR plays a role in the pathogenesis of MS, with two key markers CHOP and ATF4 found to be unregulated in the tissue of patients [77]. Further, factors associated with MS such as cytokines and oxidative stress have been associated with the upregulation of the ISR. For instance, increased expression of the cytokine IFN-*γ* in mice models eventuated in oligodendrocyte apoptosis following ISR upregulation [77]. When oligodendrocytes experience stress associated with MS, the ISR attenuates global protein synthesis so that the cell’s focus shifts from its typical protein synthesis (required for myelin production) to stress adaptation [78].

We measure stress of an oligodendrocyte by the sum of fully damaged myelin agents *σ* in its neighbourhood of 25 uniquely assigned myelin sites. We model the activation of the ISR by enforcing that any oligodendrocyte with stress level *σ* greater or equal to tolerance *ω* transitions from a myelinating to non-myelinating role. This means that once *σ* ≥ *ω* an oligodendrocyte stops repair of its 25 myelin sites. Subject to the degree of sustained activation, the ISR can remain protective (attenuated protein synthesis) or eventuate in an apoptotic event through caspase-related apoptotic signals [77, 78]. We define a subsequent tolerance *λ* that represents the one-way switch (*σ* ≥ *λ*) beyond which the oligodendrocyte undergoes apoptosis, removing the ‘health’ of its assigned myelin agents. This occurs once the oligodendrocyte is no longer myelinating, so that *ω* ≤ *λ* ≤ 25. Within the oligodendrocyte population, this gives rise to three subpopulations: myelinating, non-myelinating, and apoptotic. Given that myelinating oligodendrocytes have a half-life exceeding ten years [79], stress-induced apoptosis is their only death mechanism (see Fig 4c for further detail). Further, no new oligodendrocytes are introduced with the exception of therapeutic intervention, giving a closed population. While this neglects to account for the typical differentiation of oligodendrocyte precursor cells (OPCs) into mature oligodendrocytes, a lack of OPC differentiation in areas of lesion is thought to contribute to the progression of MS [75].

#### Myelin

To model the progressive damage and repair of myelin over the duration of the simulation, we take an approach comparable to compartmental modelling. We define a finite number S of myelin states *M_i_* where *i* ∈ {1, 2*, …,* S}. At any given time step, the state of a myelin agent represents damage events by reactivated T cells and repair events by the corresponding oligodendrocyte (see Fig 4b). The state *M_S_* represents myelin agents that are completely intact, while the state *M*_1_ represents myelin agents that are completely degraded. It follows that the intermediate states *M_i_*, where *i* ∈ {2, 3*, …,* S − 1}, represent myelin agents with intermediate degrees of damage.

It is a typical function of mature oligodendrocytes to maintain a sufficient rate of myelin turnover so that myelin replenishment and degradation are in equilibrium [80]. As such, our initial condition assumes that a population of healthy myelin is maintained by each oligodendrocyte, so that each myelin agent is initialised with state *M_S_*. Disruption of this stasis by reactivated T cells prompts transitions between the states under conditions specific to the direction of damage or repair. Demyelination occurs when a reactivated T cell occupies the same lattice site as a myelin agent. Over each time step of occupation, the myelin agent’s state updates from *M_i_*to *M_i−_*_1_. The reactivated T cell occupies the site of the myelin agent until it is fully degraded to have state *M*_1_, taking a total time of (S − 1)*τ* minutes.

Remyelination of a myelin agent requires it to be associated with a functioning, myelinating oligodendrocyte. Recall that *σ* is the sum of myelin agents with state *M*_1_ in an oligodendrocyte’s local neighbourhood of 25 myelinated lattice sites. If *σ < ω*, any compromised myelin agents in the neighbourhood with state *M_i_*, where *i* ∈ {1, 2*, …,* S − 1}, are eligible for repair by the oligodendrocyte. This reparative action only occurs in the absence of reactivated T cell presence, so that each instance of partial myelin repair from state *M_i_*to state *M_i_*_+1_ occurs after W time steps of no reactivated T cell occupation of that myelin site. The total time to repair a fully degraded myelin agent is (S − 1)W*τ* minutes, where *τ* is the time step. Note that if W = *τ* then the time taken to damage and repair myelin is equivalent.

### Model summary

We have extensively searched the literature to parameterise the model. The full set of parameters are given in the supplementary material (S1 Full parameter table). Some parameters are estimated to be within a range that we consider biologically reasonable, and are subsequently subjected to a sensitivity analysis. In Table 2 we highlight these key parameters alongside a summary of the model agents and their role in the simulation. Details about the runtime and Matlab algorithms are available in the supplementary material (S2 Model implementation and costs). Each simulation of the model is assigned a different random seed. Unless otherwise specified, all results shown in subsequent sections are the average of 40 unique simulations. Our choice to average over 40 simulations is justified in the supplementary material (S3 Simulation averaging).

**Table 2.** Summary of the five agent populations and the three key parameters identified to model treatment effects.

| Agent | Resident domain | Summary |
| --- | --- | --- |
| Primed T cell | Peripheral blood | Introduction by stochastic pulses, removal by apoptosis, conversion loss due to reactivation by perivascular macrophages. Undertakes unbiased migration and can cross BBB from peripheral blood to perivascular space. |
| Perivascular macrophage | Perivascular space | Fixed population randomly initialised in perivascular space. Undertakes unbiased migration within perivascular space and reactivates peripherally primed autoreactive T cells. |
| Reactivated T cell | Perivascular space and parenchyma | Introduction by primed T cell and PVM interaction, removal by apoptosis. Undertakes biased migration from perivascular space to parenchyma based on the presence of damaged myelin. Degrades myelin. |
| Myelin | Parenchyma | Closed, spatially fixed population. Measure of disease progression with state ranging from fully intact to fully degraded. |
| Oligodendrocyte | Parenchyma | Closed, spatially fixed population. Has three actions: myelinates, stops myelinating, and undergoes apoptosis. |
| Parameter | Description | Significance |
| $b_R$ | BBB permeability | Probability of a primed T cell crossing the BBB to enter the perivascular space from the peripheral blood. |
| $\omega$ | Remyelination threshold | Total number of damaged myelin agents belonging to an oligodendrocyte before it stops remyelinating. |
| $\lambda$ | Apoptosis threshold | Total number of damaged myelin agents belonging to an oligodendrocyte before it undergoes apoptosis. |

## Results

### ABM capabilities for typical disease course

To affirm the model’s ability to simulate a disease course typical of an untreated MS patient, we first simulated with the following parameters: *b_R_* = 0.1*, ω* = 10, and *λ* = 14. All other parameter choices follow those specified in the supplementary material (S1 Full parameter table). The average disease course that results with this parameter set is compatible with clinically observed disease progression (Fig 5). Recalling Fig 1, brain volume decreases during relapses as the heightened reactivated T cell presence (Fig 5a) contributes to myelin loss (Fig 5b). By Day 300 there is an average decrease in myelin coverage of 96% over the 3mm^2^ area of myelinated parenchyma modelled. The most significant myelin loss occurs during the first relapse event, with a diminishing rate of myelin loss with each relapse thereafter. We attribute this to the rightward progression of the lesion edge (Fig 6a), placing myelin targets further from the PVS where T cells are activated. Focused reactivated T cell migration over long distances is not promoted in this model given the bias strength is inversely proportional to the distance to the target. The myelin agents tend to be in either a fully intact or fully demyelinated state, with the intermediate states serving as transitory (Fig 5b). The profiles of oligodendrocyte agent behaviour (Fig 5c) closely follow the states of the myelin agents (Fig 5b). Similar to the intermediate myelin states, the nonmyelinating behaviour of oligodendrocytes appears to be a transitory. It is clear that the stress tolerances of *ω* = 10 and *λ* = 14 rapidly overwhelm the oligodendrocyte, with it quickly undergoing apoptosis after it stops myelinating.

**Fig 5.**
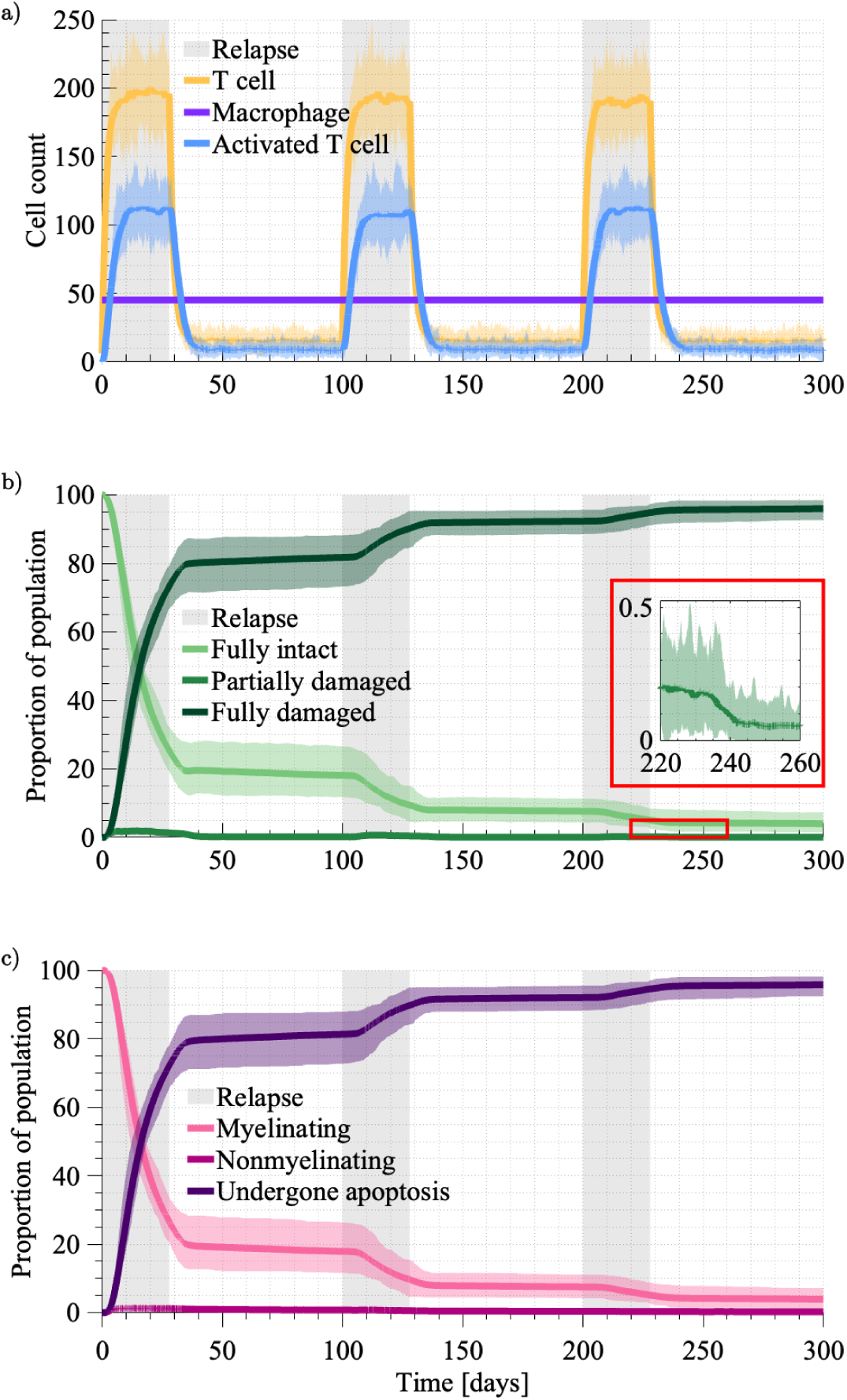
The averaged results across 40 simulations with *b_R_* = 0.1*, ω* = 10, and *λ* = 14. These results represent the expected disease course of an untreated patient. The extremes across the 40 simulations are indicated by the shading around each averaged result. The three relapse events, each 4 weeks in duration, are indicated by the grey shading. **a** The average population of primed T cells (yellow), perivascular macrophages (purple) and reactivated T cells (blue). **b** The proportion of the myelin population that is fully intact (light green), partially damaged (green) and fully damaged (dark green). **c** The proportion of oligodendrocytes that are myelinating (pink), nonmyelinating (dark pink), and have undergone apoptosis (purple).

**Fig 6.**
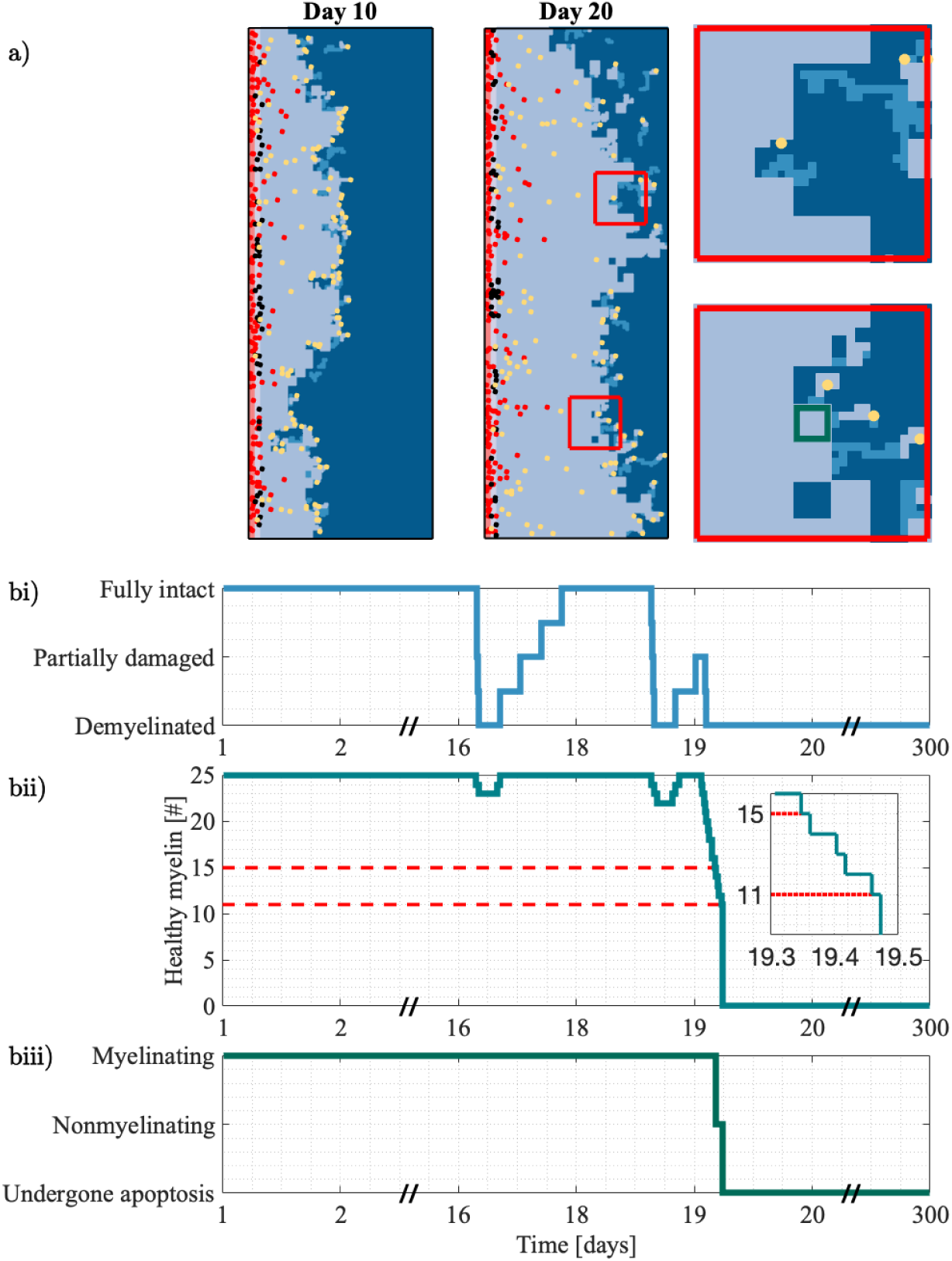
Spatial and intracellular insights from an individual simulation. **a** Shown at Day 10 and Day 20, the model produces a rightward-progressing lesion (light blue). The progression and heterogeneity of the lesion boundary are driven by the reactivated T cell population (yellow). The insets show a closeup of Day 20, where reactivated T cells create paths of intermediate damage (blue) in the densely myelinated region (dark blue). **b** The interrelationship between the level of myelin damage and the behaviour of oligodendrocytes. In **bi** we show the individual state of a myelin agent within a given oligodendrocyte (green region depicted in the inset of a). The number of healthy myelin sites within this oligodendrocyte is shown in **bii**, alongside the reductions in healthy myelin that cause the oligodendrocyte to stop myelinating and undergo apoptosis. Note that the tolerances depicted (red dashed lines) are 25-*ω* and 25-*λ* since *ω* and *λ* are given in terms of damage. The behaviour of the oligodendrocyte is shown in **biii**, where it switches from myelinating to nonmyelinating and then undergoes apoptosis once *ω* and *λ* are triggered.

In addition to averaged behaviour (Fig 5), we also considered the simulated disease course of an individual (Fig 6). The ABM provides insight at multiple spatial scales. At Day 10, reactivated T cell efforts are concentrated at the boundary, driving irregularities in the lesion edge as it forms. From the two insets, we see that modelling oligodendrocytes by 5 × 5 site areas of myelin results in block-like areas of demyelination. In regions where the oligodendrocyte is still functional, the stochasticity in each reactivated T cell’s migration is captured by paths of intermediate damage. Recalling that the functionality of individual oligodendrocytes is determined by the extent of their local myelin damage, we show the interrelationships between a selected myelin and its oligodendrocyte (Fig 6b). The individual myelin agent is degraded and repaired several times (Fig 6bi). When considering the state of all myelin agents in the neighbourhood of the oligodendrocyte, we see the demyelination of this myelin agent recorded as a reduction in the neighbourhood measure (Fig 6bii). Since the oligodendrocyte is in a myelinating state (Fig 6biii), this site and other damaged sites in the neighbourhood are remyelinated to be fully intact (Fig 6bii). Once the number of damaged myelin agents passes the tolerances *ω* = 10 and *λ* = 14 (Fig 6bii), we see the oligodendrocyte stop myelinating and undergo apoptosis (Fig 6biii). Evidently, there will be benefit in targeting oligodendrocyte resilience to increase the tolerances of *ω* and *λ*.

### Blood-brain barrier sensitivity analysis

Given the challenges in estimating the permeability of the BBB in humans, it was important to investigate the sensitivity of the model to changes in the parameter *b_R_*. To do so, we varied the BBB permeability *b_R_* from 0.005 to 0.3 and tracked the number of reactivated T cells and percentage of lost myelin. This range was informed by the expected number of T cells we expect to see in the parenchyma of an MS patient (see S1 Full parameter table). Intuitively, increasing the BBB permeability increases the number of reactivated T cells, and subsequently decreases the amount of intact myelin (Fig 7a). For most of the permeabilities simulated there is only a noticeable effect early on (Fig 7b right) after which most simulations converge to similar percentages of lost myelin (Fig 7a right). This suggests that early inflammatory events are the main driver of lesion development. To support this, prior to the second relapse there is a strong linear correlation between the maximum number of reactivated T cells and maximum percentage of lost myelin (Fig 7c first column). Following the second and third relapse events, the relationship between the two variables as a function of BBB permeability becomes less linear. This parameter clearly influences the rate at which intact myelin is lost, suggesting we can use this understanding to examine the effect of BBB-targeted DMTs. For instance, this shows it is possible to significantly reduce demyelination through reducing the BBB permeability to low enough values, such as *b_R_* = 0.005 (Fig 7a).

**Fig 7.**
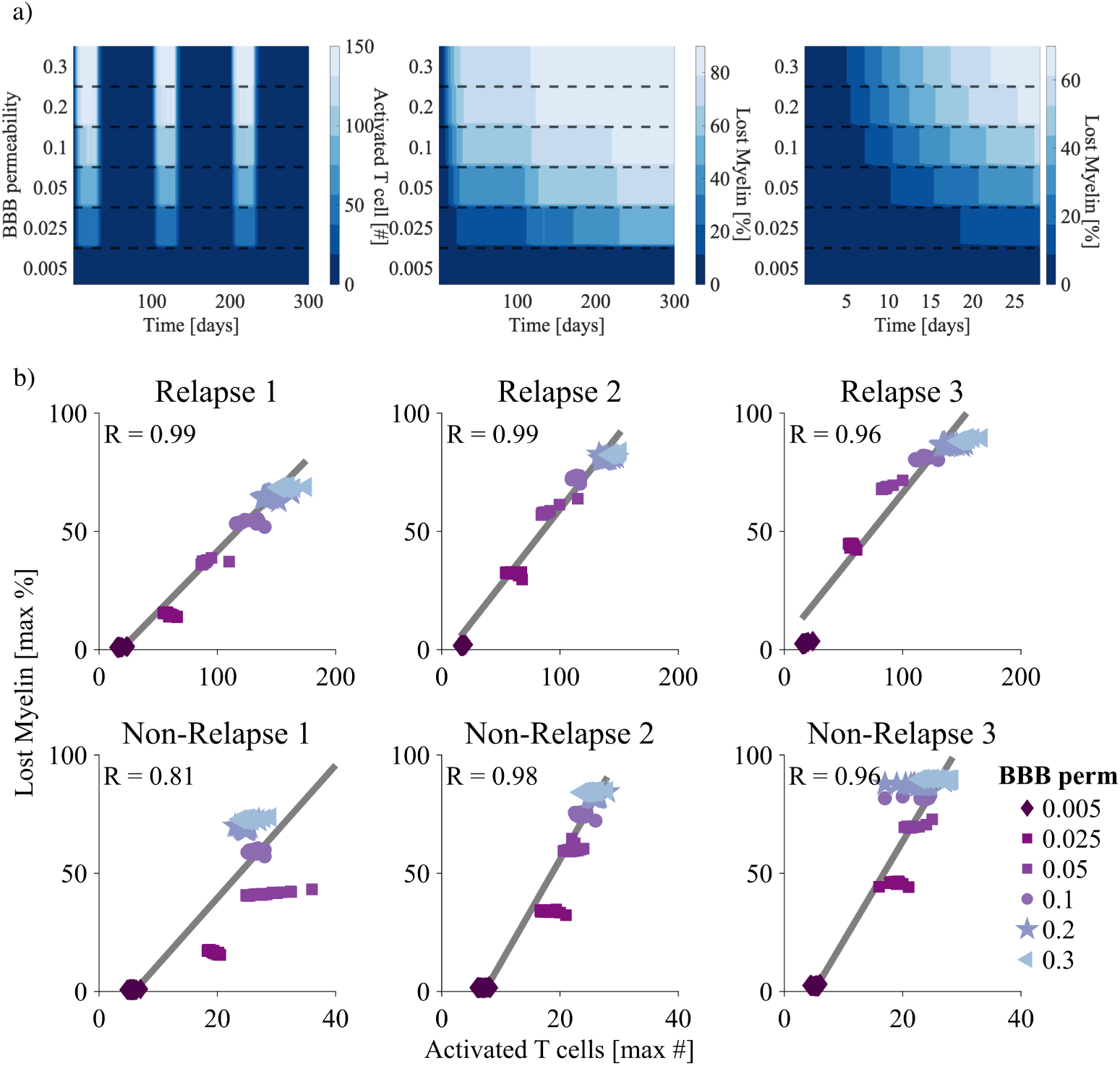
Investigation of the impact of the BBB permeability, *b_R_* on the agent-based model. **a** Heatmaps corresponding to the number of reactivated T cells (left), and percentage of lost myelin (middle and right) as a function of time (days) and BBB permeability *b_R_*. Note the timescales for the myelin heatmaps range from 300 days (middle) to 28 days (right). **b** Correlation between the number of reactivated T cells and the lost myelin percentage for the varying BBB permeabilities *b_R_* directly after each relapse (top row) and during each non-relapse period (second row). The linear correlation, *R*, is given for each.

### Oligodendrocyte sensitivity analysis

It is difficult to ascertain when the integrated stress response is upregulated in the oligodendrocytes of MS patients. As a result, we have performed a complete analysis of the model’s sensitivity to the two stress tolerances of oligodendrocytes *ω* and *λ*. Examining all possible combinations of these two parameters, where *λ* ≥ *ω*, *ω* has a strong influence of how the oligodendrocyte agents function under stress (Fig 8a). Recalling from Fig 5 that myelin states and oligodendrocyte behaviour can be conflated, significant regions of the parameter space result in rapid loss of functional oligodendrocytes, and therefore myelin. Across all combinations the nonmyelinating behaviour remains transitory, but increases when *ω* and *λ* are furthest apart.

**Fig 8.**
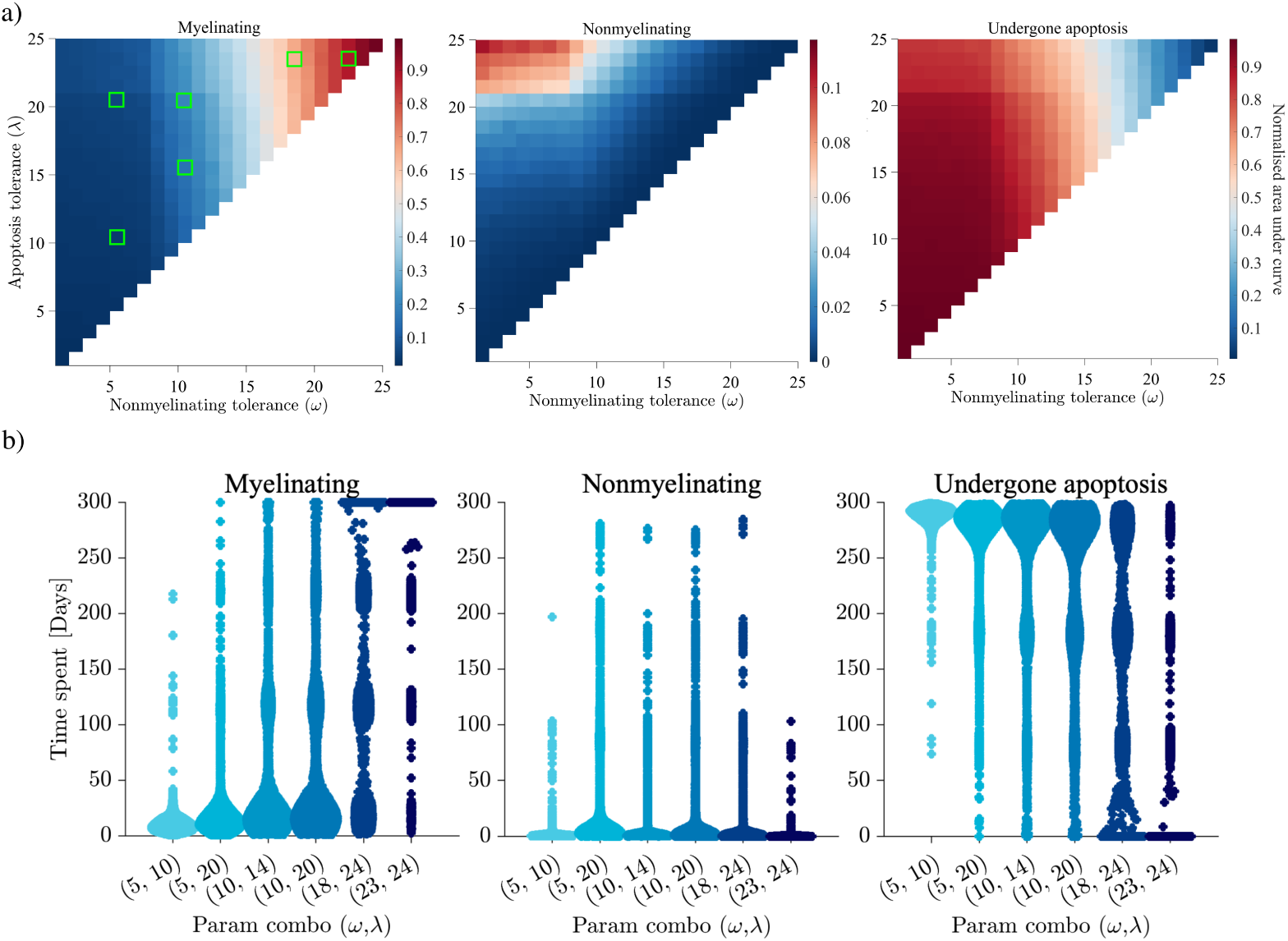
Investigation of the impact of the oligodendrocyte stress tolerances, *ω* and *λ* (recall Fig 4). **a** Heatmaps corresponding to the number of oligodendrocytes performing each behaviour (myelinating, nonmyelinating, undergone apoptosis) over the 300 day simulation. We show normalised ‘area under the curve’ data averaged across 20 simulations. The values of *ω* and *λ* range from 1-25, representing the damage required before oligodendrocytes stop myelinating and undergo apoptosis, respectively. **b** Swarm charts of select parameter pairs highlighted in (a) (green squares). We show collective data across the same 20 simulations of the frequency of oligodendrocyte time spent performing each behaviour.

For several parameter pairs (*ω*, *λ*) we highlight the time-spent by each individual performing each of the three behaviours (Fig 8b). Adverse changes in oligodendrocyte behaviour are clearly tied to relapse events, as demonstrated by the high density (increased width) of common times spent performing each behaviour. As we show collective results across 20 simulations, there is stochasticity in when these relapse-induced changes occur. For lower choices of *ω* and *λ*, oligodendrocytes are quickly overwhelmed by damage and spend the majority of the 300 day simulation having undergone apoptosis. Only when *ω* and *λ* are sufficiently high, as in the top right corner of Fig 8 (left) do oligodendrocytes retain their myelinating capabilities throughout the entire 300 day simulation. While the investigation into the BBB permeability *b_R_* showed we can control the rate of myelin loss, *ω* and *λ* appear as promising potential therapeutic targets for preventing lesion formation altogether.

### BBB-targeted therapies

As highlighted in the MS modelling work of Pennisi *et al.* (2015) [40], DMTs directly and indirectly impact the BBB to prevent its breakdown and reduce lymphocyte trafficking into the CNS. For instance natalizumab blocks the transport of T cells across the BBB by targeting their adhesion mechanisms [81]. Therapies such as glatiramer acetate and mitoxantrone instead inhibit pro-inflammatory cytokines such as IFN*γ* and TNF*α* to indirectly promote BBB preservation and regulate lymphocyte migration into the CNS [40].

When simulating the untreated disease course, we assumed that the breakdown of the BBB is invariant in space and time. However, BBB permeability reductions are a means to implicitly consider the effects of DMTs such as natalizumab. For the first treatment scenario we reduce the BBB permeability from *b_R_*= 0.1 by a factor of four to become *b_R_* = 0.025 after Day 80. In the model this is simply a reduced probability of primed T cell migration into the perivascular space from the peripheral blood, reasonably representing disrupted cell adhesion during migration processes. We compare this to the scenario where the BBB permeability has been preserved from the onset, i.e. *b_R_* = 0.025 for the whole simulation. Both scenarios are simulated with the oligodendrocyte stress tolerances of the untreated case, *ω* = 10 and *λ* = 14 (see Fig 9).

**Fig 9.**
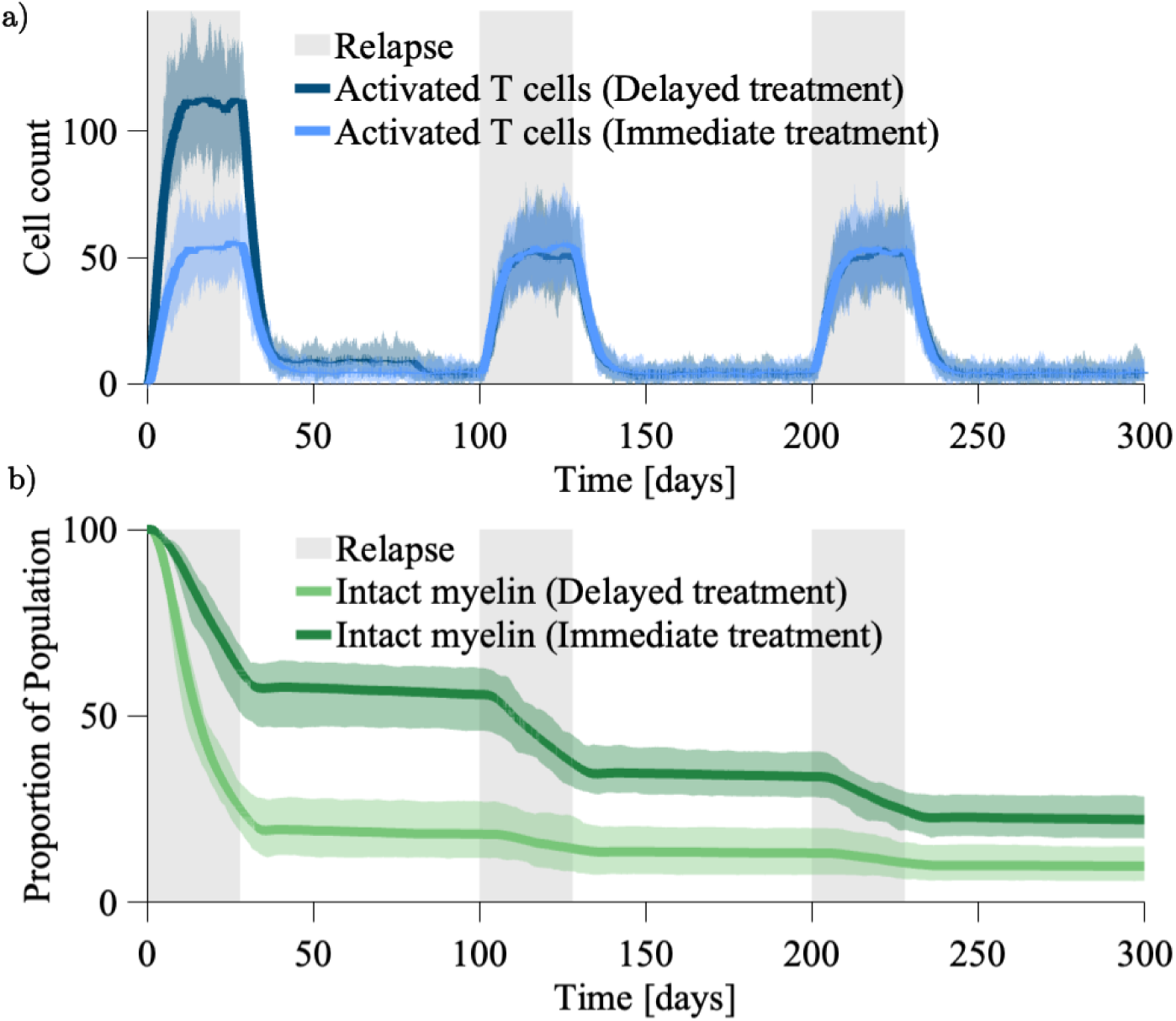
Comparison of BBB targeted treatment timing (delayed versus immediate). **a** Reactivated T cell populations under DMT treatment effects. While the typical BBB permeability is simulated as *b_R_* = 0.1, the BBB targeted treatment reduces it to *b_R_* = 0.025. The delayed treatment (dark blue) reduces the cell population from Day 80 onward, while the immediate treatment (light blue) reduces the cell population from the simulation onset. **b** The proportion of the myelin population that remains intact under the two different treatment scenarios: delayed (light green) and immediate (dark green).

As to be expected, the delayed treatment scenario (Fig 9) initially gives a reactivated T cell population of the same magnitude as the untreated case (recall Fig 5a). When we consider the patient’s proportion of intact myelin, our findings are compatible with those of Pennisi *et al.* (2015) [40]. Indeed, the best patient outcome arises in the instance that BBB opening is prevented from the simulation onset, represented by the case where *b_R_*= 0.025 for the entire simulation to give a 90.23% loss of intact myelin by Day 300. The attempt to recover the BBB integrity, represented by the case where the BBB permeability reduces from *b_R_* = 0.1 to *b_R_* = 0.025 at Day 80 sees a 77.79% loss of intact myelin by Day 300. Unfortunately, the rates of myelin loss observed in our model under these treatment strategies do not support their viability as long term solutions for preserving myelin. Both outcomes eventuate in lesion formation and significant myelin loss. The slightly improved outcome from the preventative treatment scenario relies on preservation of BBB integrity before the onset of the initial relapse event. Further, given the limited availability of DMTs for PPMS and SPMS patients this treatment strategy does not address patients with accumulated myelin loss, nor is there evidence of it promoting remyelination.

### Oligodendrocyte-targeted therapies

We now demonstrate the model’s capability for exploiting oligodendrocyte fitness as a therapeutic target. Remyelination therapies targeting oligodendrocyte retention and remyelination are not established treatments, but rather proposed in the literature or under trial. When considering exploitation of oligodendrocyte resilience, we focus on targeting mechanisms of the ISR. To simulate the effect of this therapy we consider increases in the remyelination tolerance *ω* and apoptosis tolerance *λ* to represent prolonged oligodendrocyte function and survival. Increases in the tolerances *ω* and *λ* assume oligodendrocytes to have increased stress tolerance under this treatments such as Sephin1 [27].

An additional remyelination therapy proposed in the literature is the restoration of oligodendrocytes through stem cells or OPCs [28, 29]. The proposed strategy is to restore oligodendrocyte populations in demyelinated areas for subsequent remyelination. To simulate the effects of therapy, we restore the oligodendrocyte population to be fully capable of myelinating, and the myelin population to be fully intact, at a given intervention time. This gives rise to three treatment strategies that commence at Day 80: (1) restoring oligodendrocytes (and myelin), (2) enhancing the resilience of existing oligodendrocytes through *ω* = 21, and *λ* = 24, and (3) restoring oligodendrocytes (and myelin) alongside increased resilience through *ω* = 21, and *λ* = 24. The initial relapse is simulated under the untreated parameter scheme (*b_R_* = 0.1*, ω* = 10, and *λ* = 14).

Across the three treatment strategies the profiles of myelin loss quickly diverge after intervention at Day 80 (Fig 10). The worst outcome arises when we only restore the oligodendrocyte population, i.e. treatment (1), seeing a 93.01% loss of intact myelin by Day 300 (Fig 10b). Recall that in the untreated case, there was a 96.01% loss of intact myelin by Day 300. Under the scenario where we targeted oligodendrocyte resilience, i.e. treatment (2), we see an 83.44% loss of intact myelin and stabilisation of lesion development. When we consider the combined treatment outcome we see that this is largely due to the myelin losses incurred prior to the intervention point. In the combined treatment outcome, i.e. treatment (3), we only see a 18.73% loss of intact myelin. More significantly, we observe that following the conclusion of the second and third relapses there is evidence of active remyelination, with an uptick in the intact myelin population. This is due to decreased rates of loss of our reparative cell population (oligodendrocytes). These events of repair occur once relapses diminish to an intensity that the system can handle. While the therapies targeting oligodendrocyte properties don’t give rise to lesion recovery altogether, they show dual effects of altering rates of myelin loss and enabling remyelination.

**Fig 10.**
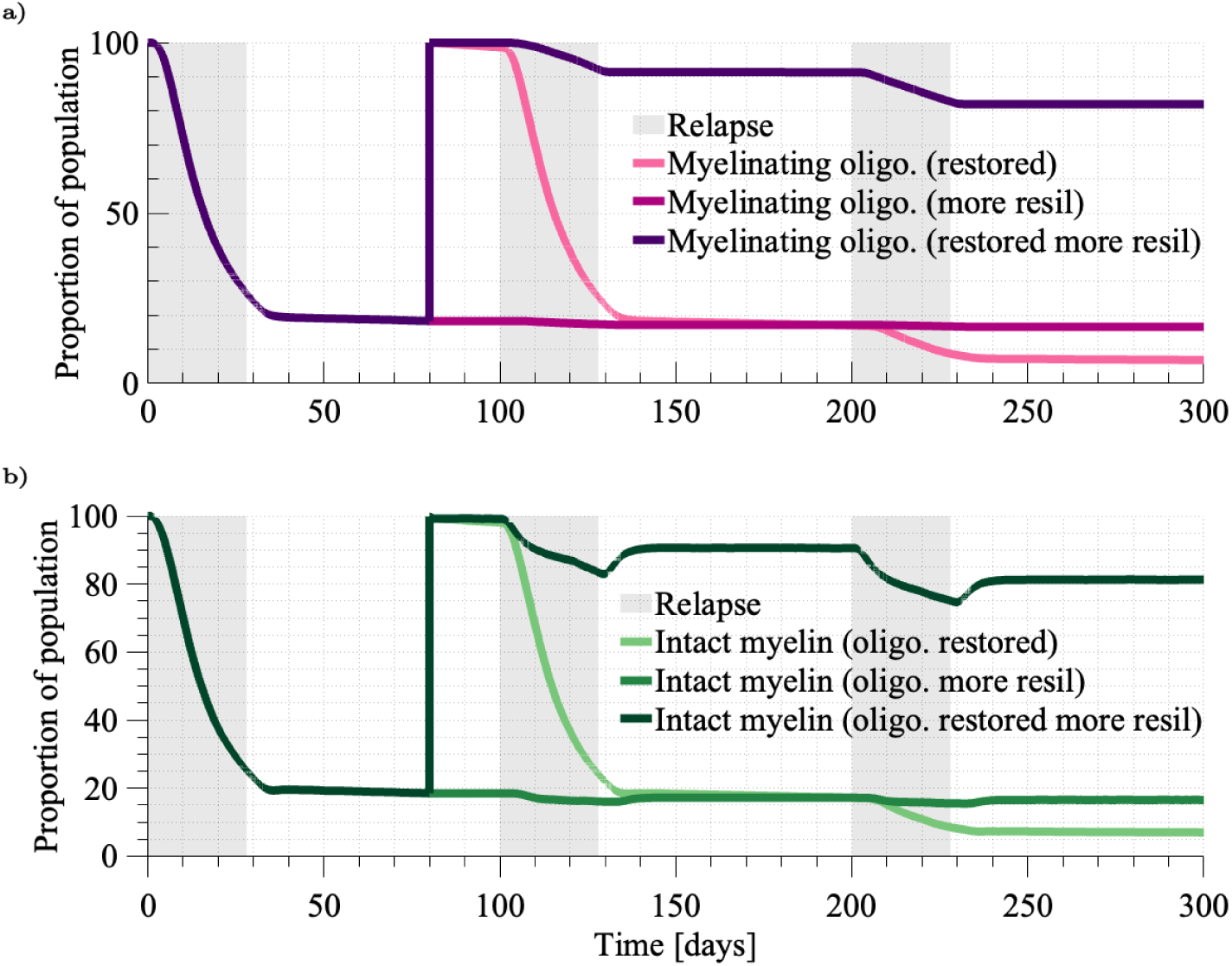
**a** Comparison of myelinating oligodendrocyte agent population across the three treatment strategies. **b** Comparison of fully intact myelin agent population across the three treatment strategies. We first simulate under the untreated parameter scheme (*b_R_* = 0.1*, ω* = 10, and *λ* = 14), and show results averaged over 40 simulations. The three treatment strategies commence at Day 80. The first strategy (1) restores oligodendrocytes (and myelin). While this restores the myelin (light green) and oligodendrocyte (light pink) populations back to 100%, they rapidly decline with the next relapses. Enhancing the resilience of existing oligodendrocytes through *ω* = 21, and *λ* = 24 under treatment (2) stabilises myelin loss (green) through the prevention of oligodendrocyte loss (dark pink). Combining these ideas in (3), we see that the intensity of the first relapse event still leads to notable myelin loss, given the abundance of potential myelin to destroy remains high.

### Combining therapeutic ideas

Given that DMTs and remyelination therapies employ different strategies for preserving myelin, we are interested in analysing the effect of combined treatment. As remyelination therapies are still in development, this combined treatment strategy exists only in the literature: Rodgers *et al*. [30] has suggested that the combined use of immunosuppressive therapies (DMTs) and remyelination therapies may be a promising strategy for improving the clinical outcomes of PPMS and SPMS patients. This is a promising prospect, given the lack of treatment available to these patients and our current inability to address their functional deficits. To simulate a combined therapeutic approach, we define an intervention point at Day 80, whereby the following parameter regime is used: *b_R_* = 0.025*, ω* = 21, and *λ* = 24. Additionally, at Day 80 we restore the parenchyma to be fully myelinated and all oligodendrocytes as myelinating. As such, all interactions from Day 80 onward can be considered as though in a patient with no existing MS-associated damage. Prior to Day 80 we simulate the untreated case (*b_R_*= 0.1*, ω* = 10, and *λ* = 14).

Under the parameter combination of *b_R_* = 0.025*, ω* = 21, and *λ* = 24 there is almost complete preservation of the fully intact myelin population at the conclusion of the 300 day simulation (Fig 11a). Interestingly, this strategy does not prevent demyelination altogether, with myelin loss occurring during the second and third relapse events.

**Fig 11.**
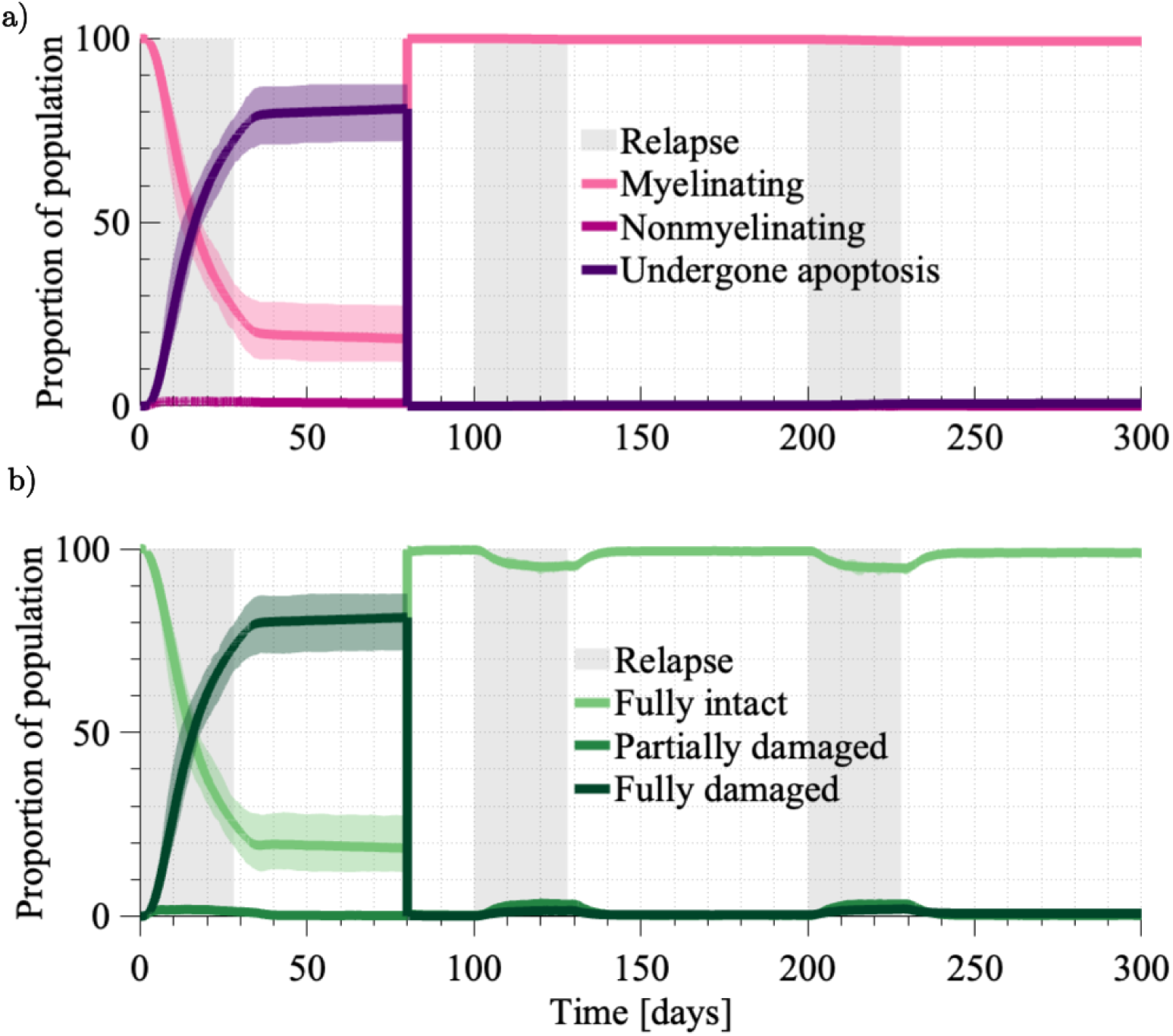
Simulation of a combined treatment approach achieving suppressed immune activity through a DMT, and the promotion of remyelination by oligodendrocyte replenishment and increases in their innate resilience. **a** Breakdown of the oligodendrocyte agent population into myelinating, nonmyelinating and having undergone apoptosis. **b** Breakdown of the myelin agent population by its level of damage (fully intact, partially damaged, fully damaged). Initially we show an untreated case where *bR* = 0.1*, ω* = 10, and *λ* = 14. Rapid myelin loss motivates the treatment intervention at Day 80, where in addition to restoring the myelin and oligodendrocyte populations we simulate under *bR* = 0.025*, ω* = 21, and *λ* = 24.

Importantly, the system is able to recover this damage. For instance, 94.5% of myelin is fully intact at Day 220, twenty days into the third relapse. By Day 300, 98.9% of the myelin population is fully intact. While we do not show the profile of the reactivated T cells, it resembles that shown in Fig 9a in the delayed DMT treatment scenario.

Excitingly, across the entire simulation there is persistent myelinating activity (Fig 11b). By Day 300, only 0.03% of the oligodendrocyte population is nonmyelinating, and only 0.7% has undergone apoptosis. This leaves most of the population capable of restoring myelin during future immune events. For this model, the system displays the ability to heal from relapse events under the increased oligodendrocyte resilience introduced by *ω* = 21 and *λ* = 24 and suppressed levels of immune activity under *b_R_* = 0.025.

Importantly, the effectiveness of this treatment strategy is not contingent on preventing demyelination altogether, but rather ensuring that myelin damage occurs at a rate that does not overwhelm local reparative ability.

## Discussion

Multiple sclerosis is a disease that presents and unfolds heterogeneously, giving rise to distinct clinical disease phenotypes. Unfortunately, patients who are not suitable for disease-modifying therapies (DMTs) see little benefit from clinical intervention besides symptomatic treatment. This has motivated discussion of a new class of treatments called remyelination therapies. Rather than suppressing disease drivers as with DMTs, these therapies are focused on enhancing a patient’s innate resilience *against* disease drivers. Target cells of these therapies are myelin-promoting cells such as oligodendrocytes. Remyelination therapies targeting oligodendrocytes remain largely unexamined clinically, and mathematical modelling of MS has typically focused on disease understanding and the effects of DMTs. In this paper, we present an agent-based model (ABM) of MS. The model is open-source and developed in Matlab, specifically designed to produce phenomena typical of MS. To our knowledge this is the first ABM to consider how immune dysregulation disrupts oligodendrocyte function on the intracellular level to impact rates of myelin loss. Given the limited clinical insight into the potential of these new therapeutic ideas, our aim is not to recapitulate existing treatment strategies but instead to generate hypotheses around the efficacy of potential strategies.

Where possible, the model was parameterised using estimates from published studies. In the instances where this was not possible we performed sensitivity analysis to understand how our parameter choices impacted the model’s reproduction of expected disease phenomena. Parameters of significant value in understanding treatment effects were the BBB permeability *b_R_*, and the tolerances beyond which oligodendrocytes stop myelinating and undergo apoptosis, *ω* and *λ* respectively. While MS models are often interested in why recurrent disease behaviour takes place [60, 82–84], we achieved relapsing-remitting dynamics by simply increasing the influx of primed T cells at predetermined times. This gave rise to three distinct relapse events with heightened disease activity. Importantly, the simulation of an untreated patient was able to produce lesion development and myelin loss typical of recurrent immune activity.

We first examined the blood-brain barrier permeability *b_R_* as a means of regulating reactivated T cell activity in the parenchyma. The BBB is both a direct and indirect target of available DMTs, with natalizumab disrupting T cell migration across the BBB [81] and glatiramer acetate and mitoxantrone acting to preserve its integrity [40]. By introducing spatial and temporal variations to BBB permeability due to pro-inflammatory cytokines, Pennisi *et al.* (2015) [40] used their ABM model to show that the prevention of BBB opening holds more potential than attempts at BBB recovery. In line with Pennisi *et al.* (2015) [40], we reduced BBB permeability *b_R_*to simulate the effect of a DMT such as natalizumab. As with their results we found that this was insufficient to stabilise lesion formation, instead only decreasing the rate of myelin loss. Based on this, we reiterate the sentiment of Pennisi *et al.* (2015) [40] that when treating with BBB-targeted DMTs for long-term myelin preservation, proactive targeting of BBB integrity is more effective than delayed treatment attempts. Once more, we highlight that the current treatments available for MS patients are disease-altering in nature, rather than restorative and able to address existing functional deficits.

More recently, oligodendrocytes have emerged as a therapeutic target capable of producing both preventative and reparative effects. The main mechanism is through the cell’s integrated stress response (ISR), prompting the development of experimental therapies such as Sephin1 that exploit this innate protective pathway [85]. Using mice with experimental autoimmune encephalomyelitis (EAE), an animal model of MS, Chen *et al.* (2019) [86] and Chen *et al.* (2021) [87] have shown Sephin1 to minimalise oligodendrocyte and myelin loss and promote remyelination. In addition to targeting oligodendrocyte properties, there is also discussion of restoring oligodendrocyte populations through stem cell or OPC-targeted therapies [28, 29]. To simulate remyelination therapies, we considered two approaches: restoration of the resident oligodendrocyte agent population, and increases in the oligodendrocyte population’s tolerances to damage, *ω* and *λ*. While the restoration and ISR exploitation strategies both improved upon the untreated disease course, only scenarios where *ω* and *λ* were increased showed clear repair of myelin following relapse events. Evidently, increased oligodendrocyte resilience was required to prevent complete overwhelm of reparative action under heightened autoreactive T cell activity. These results alone establish remyelination therapies as capable of promoting repair in MS patients.

Finally, as suggested in the literature [30], we adopted a combined treatment approach to model DMT effects through reductions in *b_R_*, alongside enhanced oligodendrocyte resilience through increases *ω* and *λ*. Under this treatment regime the system was able to maintain a steady state of high myelin coverage. It appears that the emergence of a lesion is determined by the delicate balance between the level of immune activity experienced, and the innate resilience of the oligodendrocyte population. We have shown the implication of this idea, that there are conditions that result in no lesion formation over the long term. While these results were shown following the restoration of the oligodendrocyte population following the initial relapse event, they would hold if the parameter shifts to *b_R_*, *ω*, and *λ* were applied at the onset of the simulation.

To continue the treatment investigation presented in this work, we suggest further investigation into the effects of treatment efficacy and timing. Here we assumed 100% effectiveness of the treatment strategy employed, and did not account for spatial nor temporal variations in the effectiveness the therapies. For instance, our oligodendrocyte restoration assumed all oligodendrocytes and myelin were able to be restored. While this is likely infeasible in practice, we highlight that this treatment alone did not appear to be a promising strategy given it was demonstrated to merely reset the otherwise untreated disease course under recurrent disease activity. In terms of treatment timing, our preliminary results favoured proactive treatment over delayed intervention. While this has been suggested by existing models [35, 40], this seems to be particularly true for our model given the condition preventing remyelination following oligodendrocyte loss keeps an irreversible ‘memory’ of prior immune events.

The ABM presented here has been developed specifically to capture MS and can be easily adapted to address further open questions in the MS space, of which there are numerous. We could consider local features of the vascular system, white/grey matter heterogeneity, the orientation of axons, different morphologies of the CNS and oligodendrocyte/myelin distributions, and additional immune cell populations and their respective behaviours. Many of these additions are well motivated by our results: neglecting to consider more complex vascular networks in our model imposes a limit on how far from the blood source that the lesion will progress. Further, our model assumes homogeneity within each subdomain (blood, perivascular space, parenchyma) and most existing models are similar. The ABM of Pennisi *et al*. [40] models a ‘brain’ region bordered by two blood sources each with a BBB, and the PDE system of Moise and Friedman [35] considers regions of perivascular space, demyelinated lesion, and white matter as radially-symmetric annuli surrounding a venule cross section.

Validation of the model and its potential adaptations is contingent on having a clear understanding of the lesion microenvironment, inclusive of local cellular interactions. Insights into the microenvironment of the disease may help to explain the the qualitatively distinct disease courses represented across the different patient cohorts. While the inherent stochasticity of our model introduced variability in the rates of myelin loss observed, each patient’s disease course ultimately lead to complete lesion development under the untreated parameter regime. It appears that distinct disease courses require more significant patient-to-patient variations in T cell arrival rates and distribution, BBB integrity, PVM dynamics, and oligodendrocyte resilience. Additional information at this degree of resolution about such cell distributions may be reliant on imaging modalities such as imaging mass cytometry, shown to be capable of simultaneously identifying and characterising an array of cell types in a given sample of tissue containing MS lesions [88]. Application of these imaging techniques to ABMs is already underway, with refinements to spatial statistics able to characterise cell distributions being performed using ABM data [89], and the use of imaging mass cytometry data to inform the initial conditions of agent-based, patient-specific simulations of tumour growth [90].

While the goal of reducing demyelination rates in MS patients is long-standing, there has been a recent shift in the literature regarding the strategies being employed in patient treatment. Our work is the first mathematical model of MS to consider remyelination therapies. In addition to providing an open-source framework for others to examine MS dynamics and treatments on the cellular level, our work has been able to show instances of myelin recovery under these new therapeutic ideas. While some of the therapeutic ideas we have examined are yet to be clinically examined, there are exciting experimental treatments such as Sephin1 currently under clinical trial (NCT03610334). An important implication of our work is that to prevent lesion formation, we may be able to strike a balance between acting to reduce damaging activity (DMTs/ immunosuppressants) and exploiting a patient’s innate ability to withstand it (remyelination therapies). Given the lack of curative treatment, a priority in the development of MS treatments should be understanding how to realise this balance in practice.

## Supporting information

**S1 Full parameter table.**

**S2 Model implementation and costs.**

**S3 Simulation averaging.**

## Acknowledgments

ALJ and GRW acknowledge funding from the Australian Research Council (ARC) Discovery Early Career Researcher Award (DECRA) DE240100650 and the QUT Faculty of Science Computational Bioimaging Group (CBG). ALJ also acknowledges the ARC Discovery Project (DP) DP230100025. RPA is supported by an ARC Future Fellowship (FT190100645) and also acknowledges support from ARC Discovery Project DP230100485. Computational resources used in this work were provided by the eResearch Office, Queensland University of Technology.

## References

1. Walton C, King R, Rechtman L, Kaye W, Leray E, Marrie RA, et al. Rising prevalence of multiple sclerosis worldwide: Insights from the Atlas of MS, third edition. Multiple Sclerosis (Houndmills, Basingstoke, England). 2020;26(14):1816–1821. doi:10.1177/1352458520970841.

2. Lassmann H. Axonal injury in multiple sclerosis. Journal of Neurology, Neurosurgery & Psychiatry. 2003;74(6):695–697. doi:10.1136/jnnp.74.6.695.

3. Hauser SL, Cree BAC. Treatment of Multiple Sclerosis: A Review. The American Journal of Medicine. 2020;133(12):1380–1390.e2. doi:10.1016/j.amjmed.2020.05.049.

4. Olsson T, Barcellos LF, Alfredsson L. Interactions between genetic, lifestyle and environmental risk factors for multiple sclerosis. Nature Reviews Neurology. 2017;13(11):25–36. doi:10.1038/nrneurol.2016.187.

5. Attfield KE, Jensen LT, Kaufmann M, Friese MA, Fugger L. The immunology of multiple sclerosis. Nature Reviews Immunology. 2022;doi:10.1038/s41577-022-00718-z.

6. Sospedra M, Martin R. IMMUNOLOGY OF MULTIPLE SCLEROSIS. Annual Review of Immunology. 2005;23(1):683–747. doi:10.1146/annurev.immunol.23.021704.115707.

7. Lucchinetti C, Brück W, Parisi J, Scheithauer B, Rodriguez M, Lassmann H. Heterogeneity of multiple sclerosis lesions: implications for the pathogenesis of demyelination. Annals of Neurology. 2000;47(6):707–717. doi:10.1002/1531-8249(200006)47:6¡707::aid-ana3¿3.0.co;2-q.

8. Ghasemi N, Razavi S, Nikzad E. Multiple Sclerosis: Pathogenesis, Symptoms, Diagnoses and Cell-Based Therapy. Cell Journal (Yakhteh). 2017;19(1):1–10.

9. Hemmer B, Kerschensteiner M, Korn T. Role of the innate and adaptive immune responses in the course of multiple sclerosis. The Lancet Neurology. 2015;14(4):406–419. doi:10.1016/S1474-4422(14)70305-9.

10. van Langelaar J, Rijvers L, Smolders J, van Luijn MM. B and T Cells Driving Multiple Sclerosis: Identity, Mechanisms and Potential Triggers. Frontiers in Immunology. 2020;11.

11. Zierfuss B, Larochelle C, Prat A. Blood–brain barrier dysfunction in multiple sclerosis: causes, consequences, and potential effects of therapies. The Lancet Neurology. 2024;23(1):95–109. doi:10.1016/S1474-4422(23)00377-0.

12. Teske N, Liessem A, Fischbach F, Clarner T, Beyer C, Wruck C, et al. Chemical hypoxia-induced integrated stress response activation in oligodendrocytes is mediated by the transcription factor nuclear factor (erythroid-derived 2)-like 2 (NRF2). Journal of Neurochemistry. 2018;144(3):285–301. doi:10.1111/jnc.14270.

13. Simons M, Nave KA. Oligodendrocytes: Myelination and Axonal Support. Cold Spring Harbor Perspectives in Biology. 2016;8(1):a020479. doi:10.1101/cshperspect.a020479.

14. Montalban X, Gold R, Thompson AJ, Otero-Romero S, Amato MP, Chandraratna D, et al. ECTRIMS/EAN Guideline on the pharmacological treatment of people with multiple sclerosis. Multiple Sclerosis (Houndmills, Basingstoke, England). 2018;24(2):96–120. doi:10.1177/1352458517751049.

15. Lublin FD, Reingold SC. Defining the clinical course of multiple sclerosis: results of an international survey. National Multiple Sclerosis Society (USA) Advisory Committee on Clinical Trials of New Agents in Multiple Sclerosis. Neurology. 1996;46(4):907–911. doi:10.1212/wnl.46.4.907.

16. Miller DH, Leary SM. Primary-progressive multiple sclerosis. The Lancet Neurology. 2007;6(10):903–912. doi:10.1016/S1474-4422(07)70243-0.

17. Dendrou CA, Fugger L, Friese MA. Immunopathology of multiple sclerosis. Nature Reviews Immunology. 2015;15(99):545–558. doi:10.1038/nri3871.

18. McGinley MP, Goldschmidt CH, Rae-Grant AD. Diagnosis and Treatment of Multiple Sclerosis: A Review. JAMA. 2021;325(8):765–779. doi:10.1001/jama.2020.26858.

19. Gohil K. Multiple Sclerosis: Progress, but No Cure. Pharmacy and Therapeutics. 2015;40(9):604–605.

20. Tintore M, Vidal-Jordana A, Sastre-Garriga J. Treatment of multiple sclerosis — success from bench to bedside. Nature Reviews Neurology. 2019;15(11):53–58. doi:10.1038/s41582-018-0082-z.

21. Williams T, Chataway J. Beyond ocrelizumab in primary progressive multiple sclerosis. Nature Reviews Neurology. 2022;18(1111):641–642. doi:10.1038/s41582-022-00724-8.

22. Park E, Barclay WE, Barrera A, Liao TC, Salzler HR, Reddy TE, et al. Integrin α3 promotes Th17 cell polarization and extravasation during autoimmune neuroinflammation. Science immunology. 2023;8(88):eadg7597. doi:10.1126/sciimmunol.adg7597.

23. Hellwig K, Gold R. Progressive multifocal leukoencephalopathy and natalizumab. Journal of Neurology. 2011;258(11):1920–1928. doi:10.1007/s00415-011-6116-8.

24. Olejnik P, Roszkowska Z, Adamus S, Kasarełło K. Multiple sclerosis: a narrative overview of current pharmacotherapies and emerging treatment prospects. Pharmacological Reports. 2024;doi:10.1007/s43440-024-00642-0.

25. Puniya BL, Amin R, Lichter B, Moore R, Ciurej A, Bennett SJ, et al. Integrative computational approach identifies drug targets in CD4+ T-cell-mediated immune disorders. npj Systems Biology and Applications. 2021;7(1):1–18. doi:10.1038/s41540-020-00165-3.

26. Gacem N, Nait-Oumesmar B. Oligodendrocyte Development and Regenerative Therapeutics in Multiple Sclerosis. Life. 2021;11(4):327. doi:10.3390/life11040327.

27. Chen Y, Kunjamma RB, Weiner M, Chan JR, Popko B. Prolonging the integrated stress response enhances CNS remyelination in an inflammatory environment. eLife. 2021;10:e65469. doi:10.7554/eLife.65469.

28. Franklin RJM, ffrench Constant C, Neurology PoM. Stem cell treatments and multiple sclerosis. BMJ. 2010;340:c1387. doi:10.1136/bmj.c1387.

29. Duncan I, Goldman S, Macklin W, Rao M, Weiner L, Reingold S. Stem cell therapy in multiple sclerosis: promise and controversy. Multiple Sclerosis Journal. 2008;14(4):541–546. doi:10.1177/1352458507087324.

30. Rodgers JM, Robinson AP, Miller SD. Strategies for Protecting Oligodendrocytes and Enhancing Remyelination in Multiple Sclerosis. Discovery medicine. 2013;16(86):53–63.

31. Weatherley G, Araujo RP, Dando SJ, Jenner AL. Could Mathematics be the Key to Unlocking the Mysteries of Multiple Sclerosis? Bulletin of Mathematical Biology. 2023;85(8):75. doi:10.1007/s11538-023-01181-0.

32. Khonsari RH, Calvez V. The Origins of Concentric Demyelination: Self-Organization in the Human Brain. PLOS ONE. 2007;2(1):e150. doi:10.1371/journal.pone.0000150.

33. Calvez V, Khonsari RH. Mathematical description of concentric demyelination in the human brain: Self-organization models, from Liesegang rings to chemotaxis. Mathematical and Computer Modelling. 2008;47(7–8):726–742. doi:10.1016/j.mcm.2007.06.011.

34. Lombardo MC, Barresi R, Bilotta E, Gargano F, Pantano P, Sammartino M. Demyelination patterns in a mathematical model of multiple sclerosis. Journal of Mathematical Biology. 2017;75(2):373–417. doi:10.1007/s00285-016-1087-0.

35. Moise N, Friedman A. A mathematical model of the multiple sclerosis plaque. Journal of Theoretical Biology. 2021;512:110532. doi:10.1016/j.jtbi.2020.110532.

36. Gelfand JM. Multiple sclerosis: diagnosis, differential diagnosis, and clinical presentation. Handbook of Clinical Neurology. 2014;122:269–290. doi:10.1016/B978-0-444-52001-2.00011-X.

37. Abdelgawad A, Rahayel S, Zheng YQ, Tremblay C, Vo A, Misic B, et al. Predicting longitudinal brain atrophy in Parkinson’s disease using a Susceptible-Infected-Removed agent-based model. Network Neuroscience. 2023;7(3):906–925. doi:10.1162/netna00296.

38. Sundar S, Battistoni C, McNulty R, Morales F, Gorky J, Foley H, et al. An agent-based model to investigate microbial initiation of Alzheimer’s via the olfactory system. Theoretical Biology and Medical Modelling. 2020;17(1):5. doi:10.1186/s12976-020-00123-w.

39. Weathered C, Bardehle S, Yoon C, Kumar N, Leyns CEG, Kennedy ME, et al. Microglial roles in Alzheimer’s disease: An agent-based model to elucidate microglial spatiotemporal response to beta-amyloid. CPT: pharmacometrics systems pharmacology. 2024;13(3):449–463. doi:10.1002/psp4.13095.

40. Pennisi M, Russo G, Motta S, Pappalardo F. Agent based modeling of the effects of potential treatments over the blood–brain barrier in multiple sclerosis. Journal of Immunological Methods. 2015;427:6–12. doi:10.1016/j.jim.2015.08.014.

41. Pennisi M, Rajput AM, Toldo L, Pappalardo F. Agent based modeling of Treg-Teff cross regulation in relapsing-remitting multiple sclerosis. BMC bioinformatics. 2013;14 Suppl 16(Suppl 16):S9. doi:10.1186/1471-2105-14-S16-S9.

42. Pennisi M, Russo G, Sgroi G, Palumbo GAP, Pappalardo F. In Silico Evaluation of Daclizumab and Vitamin D Effects in Multiple Sclerosis Using Agent Based Models. In: Computational Intelligence Methods for Bioinformatics and Biostatistics: 16th International Meeting, CIBB 2019, Bergamo, Italy, September 4–6, 2019, Revised Selected Papers. Berlin, Heidelberg: Springer-Verlag; 2019. p. 285–298. Available from: 10.1007/978-3-030-63061-4_25.

43. Pappalardo F, Russo G, Pennisi M, Parasiliti Palumbo GA, Sgroi G, Motta S, et al. The Potential of Computational Modeling to Predict Disease Course and Treatment Response in Patients with Relapsing Multiple Sclerosis. Cells. 2020;9(3):586. doi:10.3390/cells9030586.

44. Sips FLP, Pappalardo F, Russo G, Bursi R. In silico clinical trials for relapsing-remitting multiple sclerosis with MS TreatSim. BMC Medical Informatics and Decision Making. 2022;22(6):294. doi:10.1186/s12911-022-02034-x.

45. Júarez MA, Pennisi M, Russo G, Kiagias D, Curreli C, Viceconti M, et al. Generation of digital patients for the simulation of tuberculosis with UISS-TB. BMC Bioinformatics. 2020;21(17):449. doi:10.1186/s12859-020-03776-z.

46. Pappalardo F, Martinez Forero I, Pennisi M, Palazon A, Melero I, Motta S. SimB16: modeling induced immune system response against B16-melanoma. PloS One. 2011;6(10):e26523. doi:10.1371/journal.pone.0026523.

47. Russo G, Pennisi M, Fichera E, Motta S, Raciti G, Viceconti M, et al. In silico trial to test COVID-19 candidate vaccines: a case study with UISS platform. BMC Bioinformatics. 2020;21(17):527. doi:10.1186/s12859-020-03872-0.

48. Broome TM, Coleman RA. A mathematical model of cell death in multiple sclerosis. Journal of Neuroscience Methods. 2011;201(2):420–425. doi:10.1016/j.jneumeth.2011.08.008.

49. Cano RLE, Lopera HDE. In: Introduction to T and B lymphocytes. El Rosario University Press; 2013.Available from: https://www.ncbi.nlm.nih.gov/books/NBK459471/.

50. Wen W, Cheng J, Tang Y. Brain perivascular macrophages: current understanding and future prospects. Brain. 2024;147(1):39–55. doi:10.1093/brain/awad304.

51. Müller B, Lang S, Dominietto M, Rudin M, Schulz G, Deyhle H, et al. High-resolution tomographic imaging of microvessels. Proc SPIE. 2008;7078:70780B. doi:10.1117/12.794157.

52. Liu XY, Ma GY, Wang S, Gao Q, Guo C, Wei Q, et al. Perivascular space is associated with brain atrophy in patients with multiple sclerosis. Quantitative Imaging in Medicine and Surgery. 2022;12(2):1004–1019. doi:10.21037/qims-21-705.

53. Filippi M, Preziosa P, Banwell BL, Barkhof F, Ciccarelli O, De Stefano N, et al. Assessment of lesions on magnetic resonance imaging in multiple sclerosis: practical guidelines. Brain. 2019;142(7):1858–1875. doi:10.1093/brain/awz144.

54. Kalincik T. Multiple Sclerosis Relapses: Epidemiology, Outcomes and Management. A Systematic Review. Neuroepidemiology. 2015;44(4):199–214. doi:10.1159/000382130.

55. McDonald WI, Compston A, Edan G, Goodkin D, Hartung HP, Lublin FD, et al. Recommended diagnostic criteria for multiple sclerosis: Guidelines from the international panel on the diagnosis of multiple sclerosis. Annals of Neurology. 2001;50(1):121–127. doi:10.1002/ana.1032.

56. Cree BAC, Hollenbach JA, Bove R, Kirkish G, Sacco S, Caverzasi E, et al. Silent progression in disease activity–free relapsing multiple sclerosis. Annals of Neurology. 2019;85(5):653–666. doi:10.1002/ana.25463.

57. Vollmer T. The natural history of relapses in multiple sclerosis. Journal of the Neurological Sciences. 2007;256:S5–S13. doi:10.1016/j.jns.2007.01.065.

58. DiSano KD, Gilli F, Pachner AR. Memory B Cells in Multiple Sclerosis: Emerging Players in Disease Pathogenesis. Frontiers in Immunology. 2021;12. doi:10.3389/fimmu.2021.676686.

59. Høglund RA, Maghazachi AA. Multiple sclerosis and the role of immune cells. World Journal of Experimental Medicine. 2014;4(3):27–37. doi:10.5493/wjem.v4.i3.27.

60. Vèlez de Mendizábal N, Carneiro J, Soĺe RV, Goñi J, Bragard J, Martinez-Forero I, et al. Modeling the effector - regulatory T cell cross-regulation reveals the intrinsic character of relapses in Multiple Sclerosis. BMC Systems Biology. 2011;5(1):114. doi:10.1186/1752-0509-5-114.

61. Machado-Santos J, Saji E, Tröscher AR, Paunovic M, Liblau R, Gabriely G, et al. The compartmentalized inflammatory response in the multiple sclerosis brain is composed of tissue-resident CD8+ T lymphocytes and B cells. Brain. 2018;141(7):2066–2082. doi:10.1093/brain/awy151.

62. Karam M, Janbon H, Malkinson G, Brunet I. Heterogeneity and developmental dynamics of LYVE-1 perivascular macrophages distribution in the mouse brain. Journal of Cerebral Blood Flow Metabolism. 2022;42(10):1797–1812. doi:10.1177/0271678X221101643.

63. Kim PS, Lee PP, Levy D. A Theory of Immunodominance and Adaptive Regulation. Bulletin of Mathematical Biology. 2011;73(7):1645–1665. doi:10.1007/s11538-010-9585-5.

64. Jenner A, Yun CO, Yoon A, Coster A, Kim P. Modelling Combined Virotherapy and Immunotherapy: Strengthening the Antitumour Immune Response Mediated by IL-12 and GM-CSF Expression. Letters in Biomathematics. 2018;5(2). doi:10.30707/LiB5.2Jennera.

65. Patel AA, Ginhoux F, Yona S. Monocytes, macrophages, dendritic cells and neutrophils: an update on lifespan kinetics in health and disease. Immunology. 2021;163(3):250–261. doi:10.1111/imm.13320.

66. Bechmann I, Kwidzinski E, Kovac AD, Simbürger E, Horvath T, Gimsa U, et al. Turnover of Rat Brain Perivascular Cells. Experimental Neurology. 2001;168(2):242–249. doi:10.1006/exnr.2000.7618.

67. Krummel MF, Bartumeus F, Gèrard A. T-cell Migration, Search Strategies and Mechanisms. Nature reviews Immunology. 2016;16(3):193–201. doi:10.1038/nri.2015.16.

68. Sallusto F, Kremmer E, Palermo B, Hoy A, Ponath P, Qin S, et al. Switch in chemokine receptor expression upon TCR stimulation reveals novel homing potential for recently activated T cells. European Journal of Immunology. 1999;29(6):2037–2045. doi:10.1002/(SICI)1521-4141(199906)29:06¡2037::AID-IMMU2037¿3.0.CO;2-V.

69. Wu D. Signaling mechanisms for regulation of chemotaxis. Cell Research. 2005;15(11):52–56. doi:10.1038/sj.cr.7290265.

70. Zheng L, Guo Y, Zhai X, Zhang Y, Chen W, Zhu Z, et al. Perivascular macrophages in the CNS: From health to neurovascular diseases. CNS Neuroscience Therapeutics. 2022;28(12):1908–1920. doi:10.1111/cns.13954.

71. Huang X, Hussain B, Chang J. Peripheral inflammation and blood–brain barrier disruption: effects and mechanisms. CNS Neuroscience & Therapeutics. 2021;27(1):36–47. doi:10.1111/cns.13569.

72. Ortiz GG, Pacheco-Moiśes FP, Maćıas-Islas M Flores-Alvarado LJ, Mireles-Ramírez MA, Gonźalez-Renovato ED, et al. Role of the blood-brain barrier in multiple sclerosis. Archives of Medical Research. 2014;45(8):687–697. doi:10.1016/j.arcmed.2014.11.013.

73. Balasa R, Barcutean L, Balasa A, Motataianu A, Roman-Filip C, Manu D. The action of TH17 cells on blood brain barrier in multiple sclerosis and experimental autoimmune encephalomyelitis. Human Immunology. 2020;81(5):237–243. doi:10.1016/j.humimm.2020.02.009.

74. Miron VE, Kuhlmann T, Antel JP. Cells of the oligodendroglial lineage, myelination, and remyelination. Biochimica et Biophysica Acta (BBA) - Molecular Basis of Disease. 2011;1812(2):184–193. doi:10.1016/j.bbadis.2010.09.010.

75. Chamberlain KA, Nanescu SE, Psachoulia K, Huang JK. Oligodendrocyte regeneration: its significance in myelin replacement and neuroprotection in multiple sclerosis. Neuropharmacology. 2016;110(Pt B):633–643. doi:10.1016/j.neuropharm.2015.10.010.

76. Lin W, Kunkler PE, Harding HP, Ron D, Kraig RP, Popko B. Enhanced Integrated Stress Response Promotes Myelinating Oligodendrocyte Survival in Response to Interferon-γ. The American Journal of Pathology. 2008;173(5):1508–1517. doi:10.2353/ajpath.2008.080449.

77. Way SW, Popko B. Harnessing the integrated stress response for the treatment of multiple sclerosis. The Lancet Neurology. 2016;15(4):434–443. doi:10.1016/S1474-4422(15)00381-6.

78. Kokkosis AG, Madeira MM, Mullahy MR, Tsirka SE. Chronic stress disrupts the homeostasis and progeny progression of oligodendroglial lineage cells, associating immune oligodendrocytes with prefrontal cortex hypomyelination. Molecular Psychiatry. 2022;27(66):2833–2848. doi:10.1038/s41380-022-01512-y.

79. Tripathi RB, Jackiewicz M, McKenzie IA, Kougioumtzidou E, Grist M, Richardson WD. Remarkable Stability of Myelinating Oligodendrocytes in Mice. Cell Reports. 2017;21(2):316–323. doi:10.1016/j.celrep.2017.09.050.

80. Buscham TJ, Eichel MA, Siems SB, Werner HB. Turning to myelin turnover. Neural Regeneration Research. 2019;14(12):2063–2066. doi:10.4103/1673-5374.262569.

81. Sehr T, Proschmann U, Thomas K, Marggraf M, Straube E, Reichmann H, et al. New insights into the pharmacokinetics and pharmacodynamics of natalizumab treatment for patients with multiple sclerosis, obtained from clinical and in vitro studies. Journal of Neuroinflammation. 2016;13(1):164. doi:10.1186/s12974-016-0635-2.

82. Zhang W, Wahl LM, Yu P. Modeling and Analysis of Recurrent Autoimmune Disease. SIAM Journal on Applied Mathematics. 2014;74(6):1998–2025. doi:10.1137/140955823.

83. Zhang W, Yu P. Revealing the role of the effector-regulatory t cell loop on autoimmune disease symptoms via nonlinear analysis. Communications in Nonlinear Science and Numerical Simulation. 2021;93:105529. doi:10.1016/j.cnsns.2020.105529.

84. Dallaston MC, Birtles G, Araujo RP, Jenner AL. The effect of chemotaxis on T-cell regulatory dynamics. Journal of Mathematical Biology. 2023;87(6):84. doi:10.1007/s00285-023-02017-0.

85. Das I, Krzyzosiak A, Schneider K, Wrabetz L, D’Antonio M, Barry N, et al. Preventing proteostasis diseases by selective inhibition of a phosphatase regulatory subunit. Science. 2015;348(6231):239–242. doi:10.1126/science.aaa4484.

86. Chen Y, Podojil JR, Kunjamma RB, Jones J, Weiner M, Lin W, et al. Sephin1, which prolongs the integrated stress response, is a promising therapeutic for multiple sclerosis. Brain. 2019;142(2):344–361. doi:10.1093/brain/awy322.

87. Chen Y, Kunjamma RB, Weiner M, Chan JR, Popko B. Prolonging the integrated stress response enhances CNS remyelination in an inflammatory environment. eLife. 2021;10:e65469. doi:10.7554/eLife.65469.

88. Ramaglia V, Sheikh-Mohamed S, Legg K, Park C, Rojas OL, Zandee S, et al. Multiplexed imaging of immune cells in staged multiple sclerosis lesions by mass cytometry. eLife;8:e48051. doi:10.7554/eLife.48051.

89. Bull JA, Byrne HM. Quantification of spatial and phenotypic heterogeneity in an agent-based model of tumour-macrophage interactions. PLOS Computational Biology. 2023;19(3):e1010994. doi:10.1371/journal.pcbi.1010994.

90. Mongeon B, Hèbert-Doutreloux J, Surendran A, Karimi E, Fiset B, Quail DF, et al. Spatial computational modelling illuminates the role of the tumour microenvironment for treating glioblastoma with immunotherapies. npj Systems Biology and Applications. 2024;10(1):1–13. doi:10.1038/s41540-024-00419-4.

